# Generation and biobanking of patient-derived glioblastoma organoids and their application in CAR-T cell testing

**DOI:** 10.1101/2020.03.19.999110

**Authors:** Fadi Jacob, Guo-li Ming, Hongjun Song

## Abstract

Glioblastoma tumors exhibit extensive inter- and intra-tumoral heterogeneity, which has contributed to poor outcomes of numerous clinical trials and continues to complicate the development of effective therapeutic strategies. Current *in vitro* models do not preserve the cellular and mutational diversity of parent tumors and often require a lengthy generation time with variable efficiency. Here, we describe detailed procedures for generating glioblastoma organoids (GBOs) from surgically resected patient tumor tissue using a chemically defined medium without cell dissociation. By preserving cell-cell interactions and minimizing clonal selection, GBOs maintain the cellular heterogeneity of parent tumors. We include methods for passaging and cryopreserving GBOs for continued use, biobanking, and long-term recovery. We further describe procedures for investigating patient-specific responses to immunotherapies by co-culturing GBOs with chimeric antigen receptor (CAR) T cells. This protocol takes approximately 2-4 weeks to generate GBOs and 5-7 days to perform CAR-T cell co-culture. Competence with human cell culture, tissue processing, immunohistology, and microscopy is required for optimal results.

**Short Summary:** Detailed procedures for generating and biobanking glioblastoma organoids from resected patient tumor tissue and testing CAR-T cell efficacy by co-culture. Additional procedures for tissue processing, immunohistology, and detecting hypoxia gradients and actively proliferating cells.

**Associated Link Box for Key Reference:** Jacob, F. et al. A Patient-derived glioblastoma organoid model and biobank recapitulates inter- and intra-tumoral heterogeneity. Cell 180, 188-204 e122, doi:10.1016/j.cell.2019.11.036 (2020).

## Introduction

Tumor organoids are three-dimensional (3D) tissues generated directly from patient tumor cells that recapitulate features of their parent tumors. Recently, tumor organoids have emerged as a valuable model for many cancers, including bladder, breast, gastrointestinal, liver, ovarian, pancreatic, prostate, and rectal cancers^1-8^. These organoid models and biobanks have furthered our understanding of tumor biology and shown promise for testing new therapeutic strategies^9,10^. We recently developed a protocol to generate glioblastoma organoids (GBOs) from surgically resected patient glioblastoma tissue with minimal perturbation^11^. We showed that GBOs recapitulate the heterogeneity of their parent tumors as evidenced by immunohistology, bulk and single-cell transcriptomics, and mutational analyses revealing the maintenance of diverse cell populations and their gene expression and mutation profiles. GBOs exhibit rapid, aggressive infiltration of tumor cells when transplanted into adult rodent brains, a defining feature of human glioblastoma. We demonstrated the utility of GBOs to test personalized therapies by correlating GBO mutation profiles with responses to specific drugs and developed methods to co-culture GBOs with CAR-T cells to test killing efficacy and target specificity^11^. To better enable adoption of our methods in other laboratories, here we describe detailed procedures for generating, culturing, passaging, cryopreserving, and thawing GBOs (Figs. 1, 2). We include optimized protocols for processing tumor tissue and GBOs for downstream analyses (Box 1), performing immunohistology (Box 2, Fig. 3), and detecting hypoxia gradients and actively proliferating cells in GBOs (Box 3, Fig. 4). Furthermore, we describe procedures for co-culturing GBOs with CAR-T cells and analyzing target antigen loss, tumor cell killing, and T cell invasion, proliferation, and cytokine secretion (Fig. 5).

**Figure 1.**
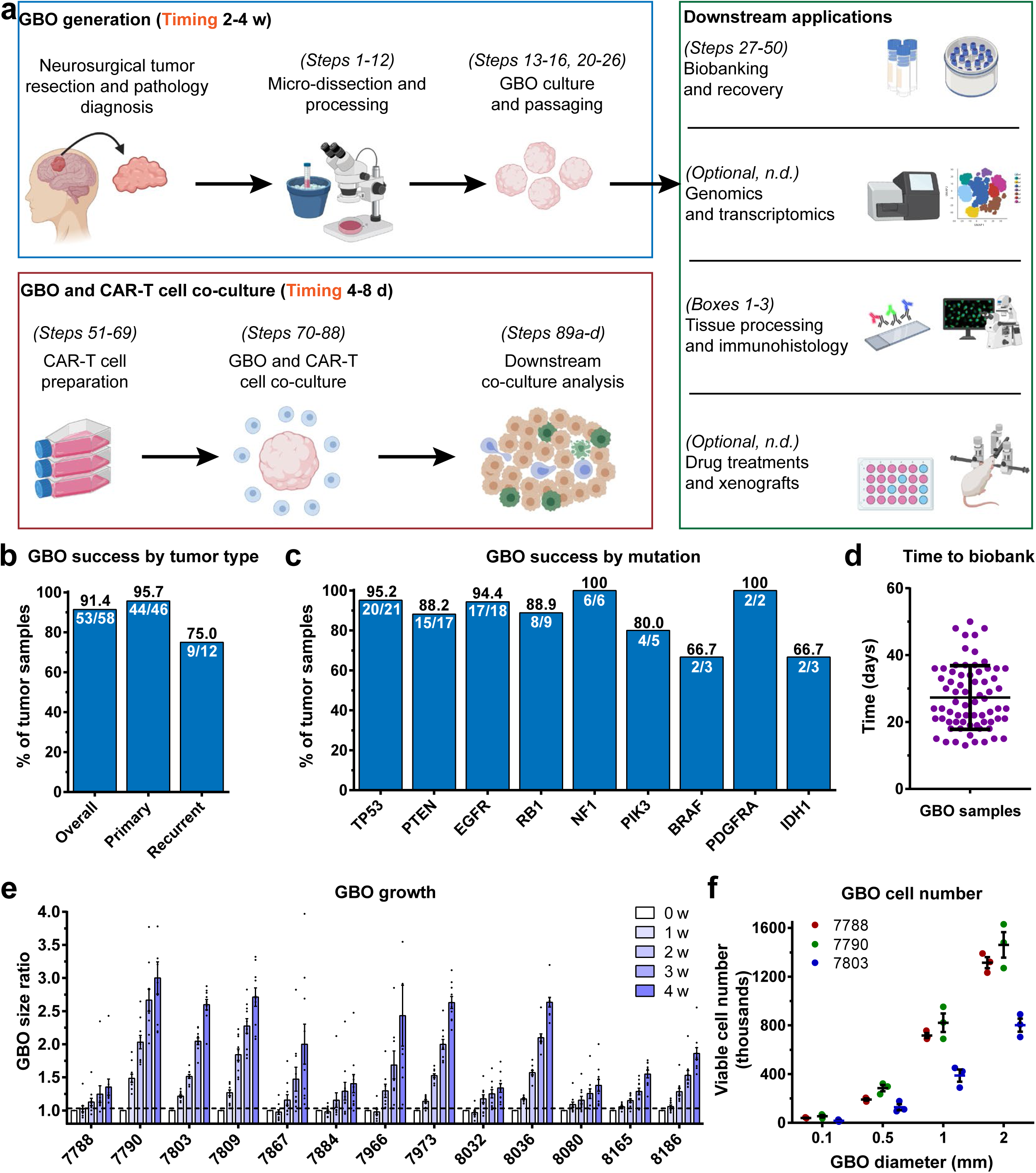
Overview of the procedures to generate GBOs from resected tumor tissue and co-culture GBOs with CAR-T cells. (a) Diagram outlining the major steps to generate GBOs from resected tumor tissue with downstream applications and to prepare and co-culture CAR-T cells with GBOs. Procedures for genomic and transcriptomic analyses, drug treatments, and xenografts are not described (n.d.) in this protocol; refer to Jacob et al., 2020 for methods and sample data for these procedures^11^. Panel created with images from www.biorender.com. (b-c) Quantification of the success rate for generating GBOs from primary and recurrent/residual tumor tissue (b) and for generating GBOs from tumors containing mutations commonly found in glioblastomas^55^ (c). The GBO success percentage is indicated in black text above each bar and the fraction of successful samples over the total samples is indicated in white text below each bar. Data plotted from information in Table S1 from Jacob et al., 2020^11^. (d) Quantification of the time from the initial tumor resection to establishment of a GBO biobank for 70 distinct GBO samples. Individual values are plotted with the mean ± SD. (e) Quantification of GBO growth over time by calculating the ratio of the measured 2D area at each time point to the 2D area at time point 0 for individual GBOs recovered from the biobank. Values represent mean ± SEM (n = 10 GBOs per sample). Data replotted from Figure 2F in Jacob et al., 2020^11^. (f) Quantification of the viable cell number within GBOs generated from three separate tumors at different GBO diameters. Values represent data from individual GBOs with the mean ± SEM (n = 3 GBOs per sample).

**Figure 2.**
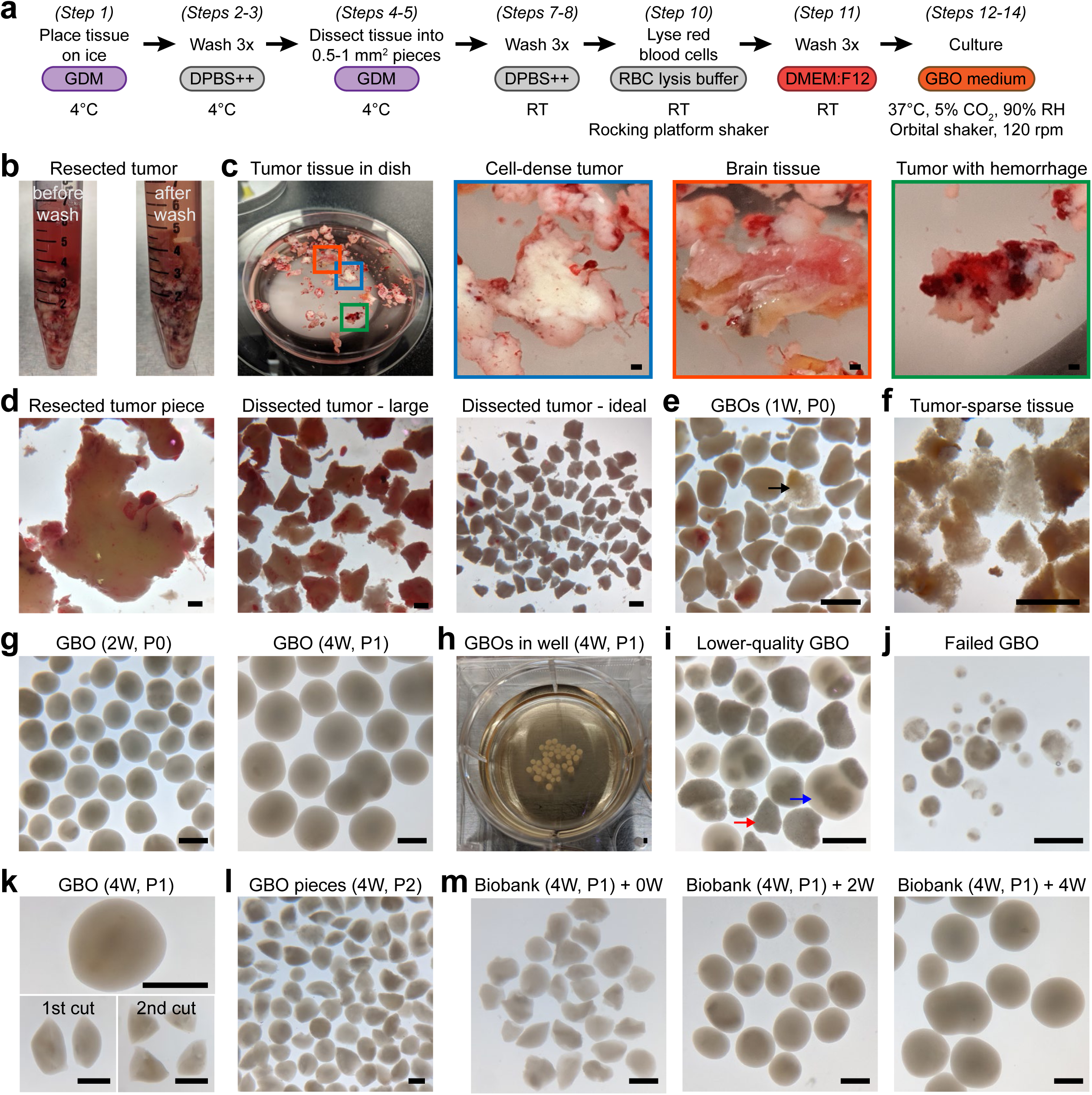
Establishment and maintenance of GBO culture. (a) Diagram outlining the steps to process glioblastoma tissue into tumor pieces for GBO culture with the medium and culture conditions indicated below each step. GDM, glioblastoma dissection medium; RBC, red blood cell; RT, room temperature; RH, relative humidity. (b) Sample bright-field images of conical tubes containing example glioblastoma tissue received from surgical resection before washing (left) and after washing and adding glioblastoma dissection medium (right). (c) Sample bright-field image of resected tumor tissue in a glass dissection dish (left) with higher magnification images of boxed regions on the right showing examples of cell-dense tumor pieces (blue box), brain tissue (red box), and tumor tissue with substantial hemorrhage (green box). Scale bars, 1 mm. (d) Sample bright-field images of a cell-dense resected tumor piece (left), large dissected tumor pieces (middle), and ideally sized dissected tumor pieces (right) after micro-dissection with fine spring dissection scissors. Scale bars, 1 mm. (e-m) Sample bright-field images of tumor pieces in culture after 1 week (e; black arrow indicates tumor-sparse tissue), tumor-sparse tissue in culture after 1 week that should be removed (f), GBOs after 2 weeks (g; left) and 4 weeks in culture (g; right), GBOs in a 6-well culture plate after 4 weeks of culture showing the approximate GBO size when ready to passage (h), lower-quality GBOs after 4 weeks in culture that may be removed (i; red arrow indicates tumor-sparse necrotic tissue and blue arrow indicates cell-dense tumor tissue growing outwards from tumor-sparse necrotic tissue), example of failed GBOs after 4 weeks in culture (j), a whole GBO after 4 weeks in culture before passaging (k; top) and after 1^st^ (k; bottom-left) and 2^nd^ (k; bottom-right) bisection, GBO pieces immediately after passaging (l), and GBOs that were cryopreserved after 4 weeks in culture and thawed from the biobank (m; left) and cultured for 2 weeks (m; middle) and 4 weeks (m; right). The age in weeks (W) and the passage number (P) of the GBOs are indicated in the parenthesis after each label. Scale bars, 1 mm.

**Figure 3.**
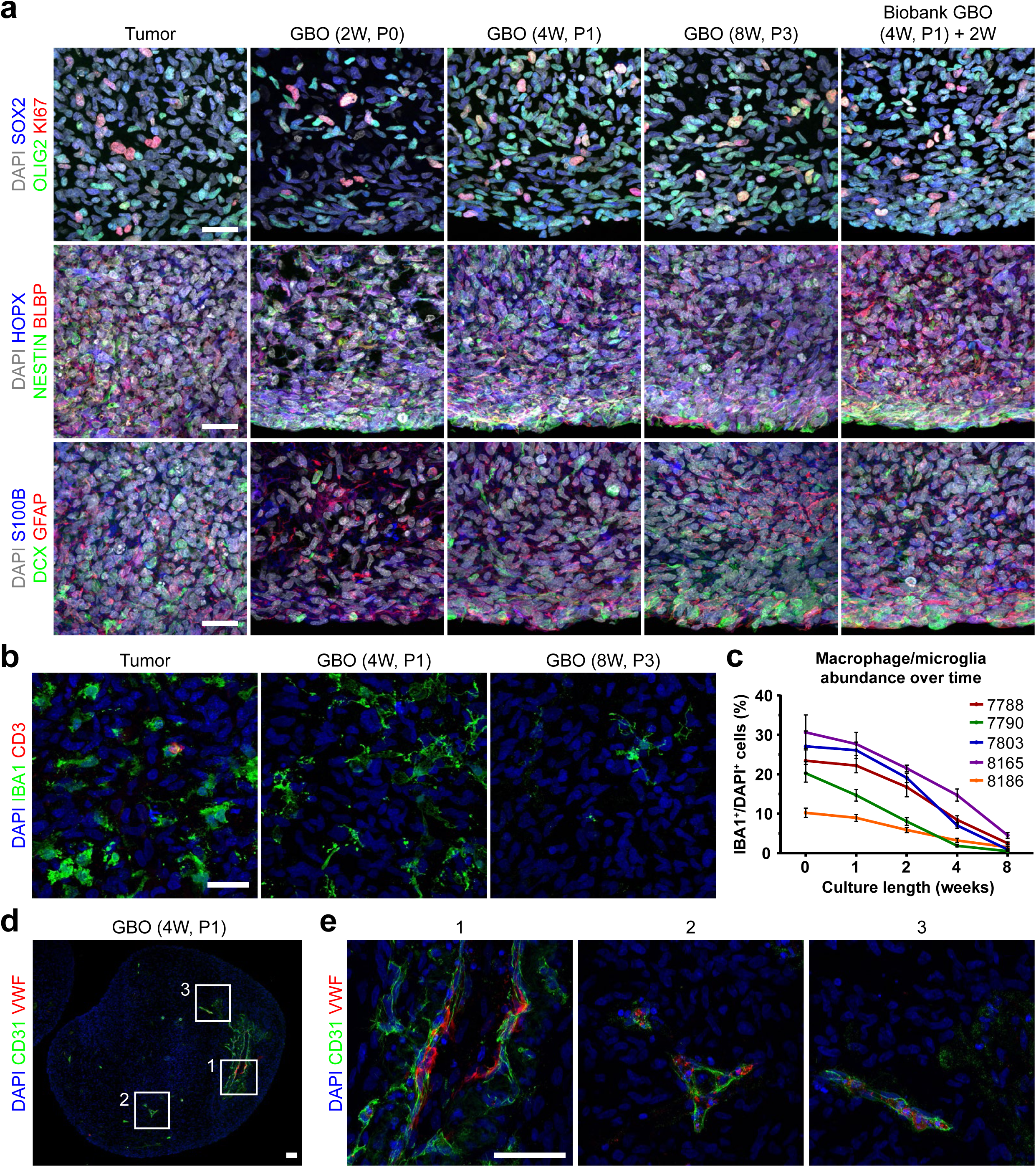
Immunofluorescence characterization of GBOs and parent tumors. (a) Sample confocal images of fluorescent immunohistology for different markers showing the maintenance of parent tumor marker expression, cellular composition, and proliferation in cultured GBOs at different ages and GBOs thawed from the biobank and cultured for 2 weeks for one patient. Representative images are based on the analysis of 5 tumor pieces or GBOs per time point. The age in weeks (W) and the passage number (P) of the GBOs are indicated in the parenthesis after each label. Fluorescent immunohistology and marker quantification for several GBO samples at multiple time points can be found in Figures 2A-B, S1A-B, and S2D of Jacob et al., 2020^11^. Scale bars, 25 μm. (b) Sample confocal images of fluorescent immunohistology for macrophage/microglia marker IBA1 and T cell marker CD3 in the parent tumor and corresponding GBOs at 4 weeks and 8 weeks showing the maintenance, but gradual loss of immune cells. Representative images are based on the analysis of 5 tumor pieces or GBOs per time point. Scale bar, 25 μm. (c) Quantification of the percentage of IBA1^+^/DAPI^+^ macrophage/microglia cells in GBOs at different ages for 5 GBO samples. Values represent mean ± SEM (n = 5 GBOs per sample and time point with 3 images per GBO). (d) Sample confocal image of fluorescent immunohistology for endothelial cell markers CD31 and Von Willebrand factor (VWF) in a GBO at 4 weeks showing maintenance of some vasculature in GBOs. Representative image is based on the analysis of 5 GBOs. Scale bar, 50 μm. (e) Higher magnification confocal images of boxed regions in (d) with fluorescent immunohistology for CD31 and VWF showing variations in vessel morphology. Scale bar, 50 μm.

**Figure 4.**
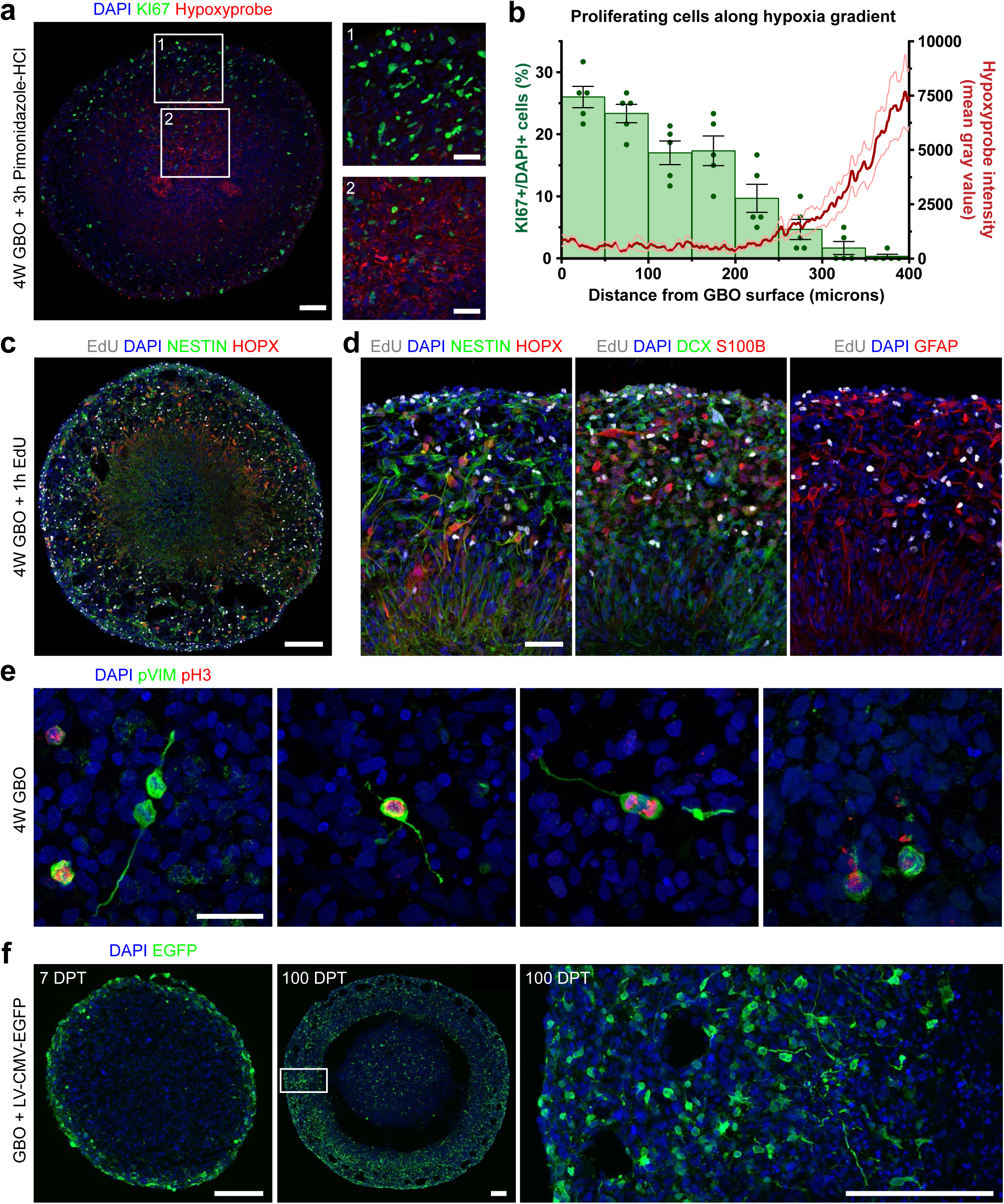
Detection of hypoxia gradients and actively proliferating cells in GBOs. (a) Sample confocal images of fluorescent immunohistology for pimonidazole-HCl (Hypoxyprobe) and KI67 showing the hypoxia gradient present in a large GBO (left) with higher magnification images of boxed regions highlighting differences in the number of KI67^+^ cells in the GBO periphery (right-top) and core (right-bottom). Representative image is based on the analysis of 5 GBOs. Scale bars, 100 μm (left) and 50 μm (right). (b) Quantifications of Hypoxyprobe fluorescent intensity as a function of distance from the surface to the center of GBOs (red) and KI67^+^ cell distance from the surface of GBOs (green) showing fewer KI67^+^ cells located within the more hypoxic regions at the center of GBOs. Values for Hypoxyprobe fluorescent intensity represent the mean ± SEM (n = 5 GBOs from a single sample) and values for KI67^+^ cell distance are represented as a histogram with the mean ± SEM with individual values plotted for each 50 micron interval (n = 5 GBOs from a single sample). (c) Sample confocal image of fluorescent immunohistology in a large GBO showing EdU detection in tumor cells labeled with NESTIN and HOPX after a 1 hour incubation with 1 μM EdU. Representative images are based on the analysis of 5 GBOs. Scale bar, 200 μm. (d) Higher magnification sample confocal images of fluorescent immunohistology in a large GBO showing EdU detection in tumor cells labeled with NESTIN, HOPX, DCX, S100B, and/or GFAP after a 1 hour incubation with 1 μM EdU. Images highlight many radially oriented cells located outside the core. Representative images are based on the analysis of 5 GBOs. Scale bar, 50 μm. (e) Sample confocal images of fluorescent immunohistology for phospho-vimentin (pVIM) and phospho-histone H3 (PH3) in GBOs cultured for 4 weeks highlighting the diverse morphology of tumor cells in mitosis. Representative images are based on the analysis of 5 GBOs. Scale bar, 25 μm. (f) Sample confocal images of fluorescent immunohistology for EGFP in GBOs transduced with CMV-EGFP lentivirus for 2 hours and cultured for 7 days (left) and 100 days (middle) without passaging to label tumor cells and their progeny. Lentivirus initially efficiently infects cells on the surface of GBOs that generate many labeled progenies distributed throughout the GBOs after extended culture. A higher magnification image (right) highlights the diverse morphology of labeled cells after 100 days post transduction (DPT). Representative images are based on the analysis of 5 GBOs per timepoint. Scale bars, 200 μm.

**Figure 5.**
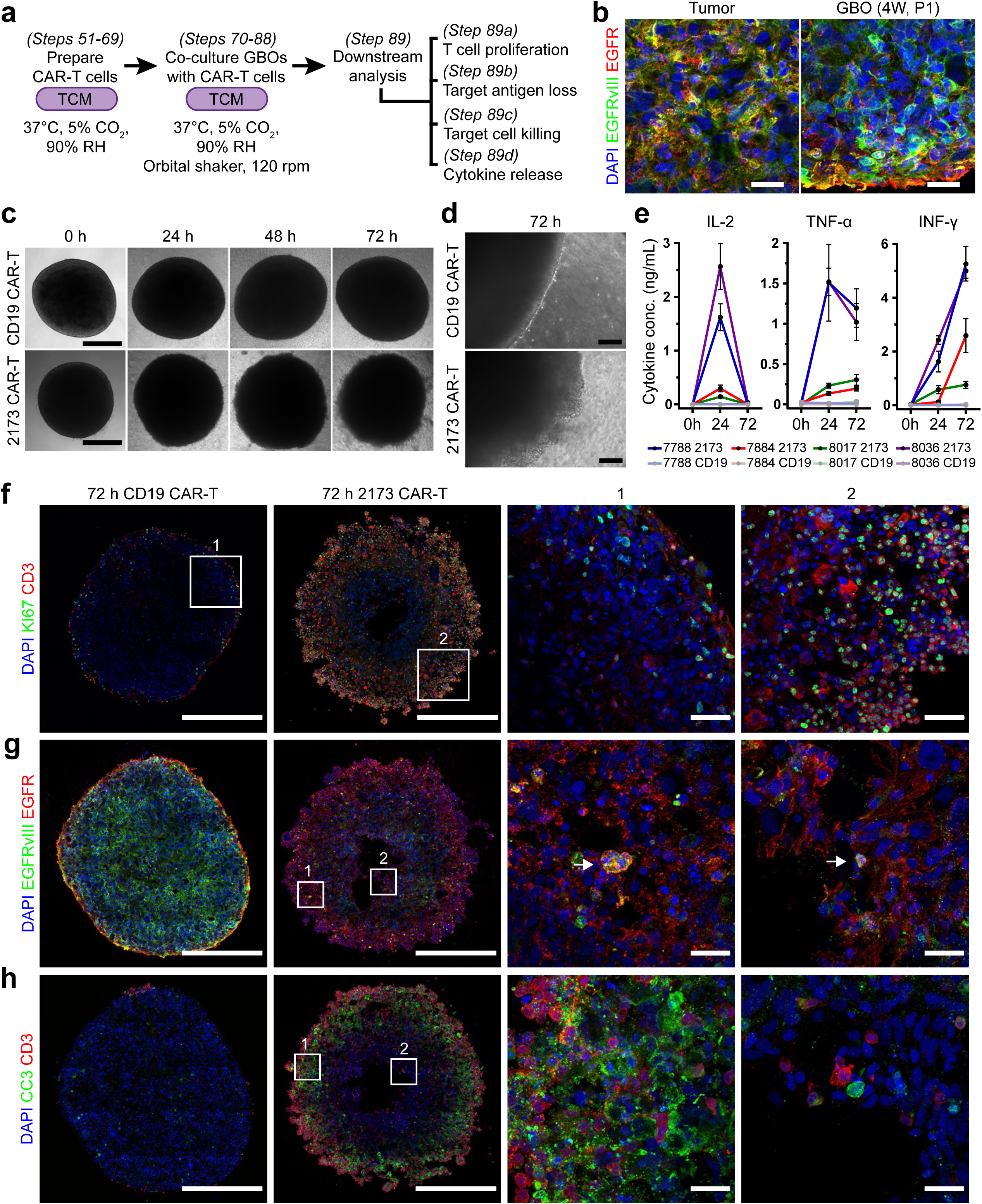
GBO co-culture with CAR-T cells. (a) Diagram outlining the steps to prepare and co-culture CAR-T cells with GBOs with downstream analyses. The medium and culture conditions are indicated below each step. TCM, T cell medium; RH, relative humidity. (b) Sample confocal images of fluorescent immunohistology for EGFR and EGFRvIII in parent tumor tissue and corresponding GBOs showing the maintenance of heterogeneous EGFRvIII expression. Representative images are based on the analysis of 5 tumor pieces or GBOs. Scale bar, 25 μm. (c) Sample bright-field images of individual EGFRvIII-expressing GBOs monitored at 0, 24, 48, and 72 hour time points when co-cultured with either CD19 or 2173 CAR-T cells. Representative images are based on the analysis of 10 GBOs. Scale bar, 500 μm. (d) Sample bright-field images of the edge of EGFRvIII-expressing GBOs when co-cultured with either CD19 or 2173 CAR-T cells for 72 hours highlighting differences in T cell invasion and proliferation. Representative images are based on the analysis of 10 GBOs. Scale bar, 100 μm. (e) Quantification of IL-2, TNF-α, and IFN-γ concentrations in media collected at 0, 24, and 72 hours after co-culture of GBOs with either CD19 or 2173BBz CAR-T cells by ELISA. Values represent mean ± SEM (n = 3 GBOs per sample per timepoint). Data is replotted from Figure S7D in Jacob et al., 2020^11^. (f) Sample confocal images of fluorescent immunohistology for CD3 and KI67 in EGFRvIII-expressing GBOs after co-culture with either CD19 or 2173 CAR-T cells for 72 hours showing increased T cell invasion and proliferation when co-cultured with 2173 CAR-T cells with higher magnification images of boxed regions highlighting these differences. Representative images are based on the analysis of 5 GBOs. Scale bars, 500 μm (whole GBO) and 50 μm (boxed regions). (g) Sample confocal images of fluorescent immunohistology for EGFRvIII and EGFR in EGFRvIII-expressing GBOs after co-culture with either CD19 or 2173 CAR-T cells for 72 hours showing EGFRvIII antigen loss when co-cultured with 2173 CAR-T cells with higher magnification images of boxed regions highlighting loss of EGFRvIII antigen at the GBO periphery (1) and core (2). White arrows highlight EGFRvIII^+^ cells with karyohexis. Representative images are based on the analysis of 5 GBOs. Scale bars, 500 μm (whole GBO) and 25 μm (boxed regions). (h) Sample confocal images of fluorescent immunohistology for cleaved caspase 3 (CC3) and CD3 in EGFRvIII-expressing GBOs after co-culture with either CD19 or 2173 CAR-T cells for 72 hours showing increased apoptosis when co-cultured with 2173 CAR-T cells with higher magnification images of boxed regions highlighting apoptotic cells near T cells at the GBO periphery (1) and core (2). Representative images are based on the analysis of 5 GBOs. Scale bars, 500 μm (whole GBO) and 25 μm (boxed regions).

### Development of glioblastoma organoid culture and biobanking methodology

An ideal *in vitro* model for glioblastoma would maintain cellular diversity, preserve native cell-cell interactions, recapitulate defining histological characteristics, and maintain the transcriptomes and mutation profiles of parent tumors. To be useful for personalized medicine approaches, the model would need a quick generation time with high fidelity for success and be scalable to test potential therapeutics. This would require maintaining patient-derived glioblastoma tissue in culture with minimal perturbation. Towards this goal, we opted to directly culture micro-dissected tumor pieces without cell dissociation to preserve cell-cell interactions and minimize clonal selection. We start from resected patient tumor tissue that can be sampled from different tumor foci or subdivided into regional samples to analyze intra-tumoral heterogeneity. Through testing of many medium formulations, we learned that tumor pieces did not require basic fibroblast growth factor (bFGF), epidermal growth factor (EGF), serum, or added extracellular matrix such as Matrigel to survive, form organoids, and proliferate at rates similar to parent tumors^11^. We optimized a chemically defined medium that contains basic nutrients and components needed to support glial cell health with few exogenous factors to minimize clonal selection induced by growth factors and decrease potential treatment confounds. This resulted in an optimized version of our medium used to maintain human brain organoid cultures^12,13^. We culture tumor pieces on an orbital shaker to increase oxygen and nutrient diffusion and aid the formation of spherical organoids. Because of these conditions, GBOs can grow quite large, recapitulating hypoxia gradients and cell distributions found in glioblastoma tumors^11^. GBOs are generated in 2-4 weeks, making them useful for therapeutic testing on a clinically relevant timescale. To extend the usefulness of GBOs, we developed optimized procedures to cryopreserve 3D tissue that has been traditionally difficult to biobank owing to poor penetration of DMSO into cells within the core. We address this problem by cutting GBOs into small pieces and pre-incubating them in DMSO-containing freezing medium to allow complete perfusion before freezing. We also incubate GBO pieces with Y-27632 (ROCK inhibitor) before freezing and during thawing to help inhibit cell death^14-17^. This methodology has enabled the establishment of a living biobank of GBOs for future research.

### Development of CAR-T cell co-culture methodology

Although CAR-T cell therapy has shown success in several blood cancers^18^, it has not been as effective in solid tumors^19^. Current *in vitro* models for solid tumors often lack cellular heterogeneity and do not maintain the mutations that lead to altered surface antigens, such as the EGFRvIII variant in glioblastomas which has shown unclear efficacy in clinical trials using CAR-T cells that recognize EGFRvIII specifically^20,21^. Because of these constraints, experiments to test the efficacy of CAR-T cells have often relied on overexpression of target antigens in tumor cell lines, which may not recapitulate the endogenous condition^22^. Because of their three-dimensional architecture, tumor organoids have recently been used as a model to test CAR-T cell killing^23^. To mimic CAR-T cell invasion into solid tumors, such as glioblastoma, we developed an optimized method to co-culture CAR-T cells with GBOs. Because GBOs maintain the cellular heterogeneity and cell-cell interactions of parent tumors, they better recapitulate the native tumor environment compared to adherent or suspension cell models. GBOs also maintain the endogenous expression of mutant proteins, such as EGFRvIII, which has been traditionally difficult to maintain in cell lines. We used the recently developed 2173-BBz anti-EGFRvIII CAR-T cell that specifically binds mutant EGFRvIII commonly found in glioblastomas^21^ to show the usefulness of GBOs to test the efficacy and specificity of CAR-T cells when co-cultured. We optimized a ratio of CAR-T cells to tumor cells to model and observe CAR-T cell infiltration and proliferation, target antigen loss, and tumor cell killing on a clinically relevant timescale. Using these methods, the efficacy and specificity of engineered CAR-T cells can be tested *in vitro* before initiating therapy in patients to stratify patients for clinical trials and to better predict therapeutic responses.

### Applications of GBOs to study glioblastoma biology and test therapeutics

Our protocol provides a 3D *in vitro* model for glioblastoma that is generated directly from resected patient tumor tissue with minimal perturbation and quick generation time. Our model allows one to study glioblastoma biology in real time and provides a platform to test targeted therapeutics, such as radiation, drugs, or CAR-T cells for personalized medicine^11^. The maintenance of parent tumor cellular heterogeneity and key driver mutations makes our GBO model useful for studying glioblastoma progenitor subtypes and lineage hierarchies. The presence of different tumor cell types or cellular states can be studied by marker immunohistology and single-cell RNA sequencing, similar to what has been done with patient tissue^24,25^. Single-cell sequencing can be used to identify putative cancer stem and progenitor cells and infer their lineage trajectories. GBOs are readily amenable to treatment with nucleoside analogs (e.g. EdU) or viruses to label cell populations (Fig. 4c, d, f). EdU pulse-chase experiments can be used to study population lineage dynamics^11^ and lentiviruses and retroviruses could be used to label progenitors and trace their progeny by permanent fluorescent labeling or RNA barcoding^26^. GBOs may also be subjected to genetic manipulations, such as overexpression, shRNA/siRNA knockdowns and CRISPR-mediated knockouts to study genetic contributions to the disease. The dynamic behavior of glioblastoma cells, such as cell division and migration can be studied by two-photon or confocal-based live imaging of fluorescently labeled tumor cells. Additionally, cell types and mechanisms underlying tumor cell infiltration into the surrounding brain tissue can be studied by orthotopic xenograft of GBOs into rodent brains^11^.

Our procedure for generating glioblastoma organoids is advantageous because GBOs can be generated in as little as 2-4 weeks following surgical resection. This quick generation time allows for testing of targeted therapies *in vitro* before the initiation of treatment in patients. Data gathered from immunohistology, transcriptomics, and genomics could inform the design of drug panels whose efficacy would be tested in GBOs. By maintaining cultures after drug and/or radiation treatment, GBOs provide a system to study which tumor populations can resist and expand after treatment. Continued culture after drug withdrawal may provide insights on which cell populations lead to tumor recurrence. The efficacy of CAR-T cells which recognize different target antigens can be tested in isolation or in combination by co-culture with GBOs to develop new therapies for clinical trials. Our protocol for CAR-T cell co-culture can also be applied to test CAR-T cells for other tumor organoids, although the conditions may need optimization for each tumor type. GBOs also offer a valuable resource to identify novel targetable surface antigens by incubation with naïve peripheral T cells, similar to what has been reported for colorectal cancer and non-small-cell lung cancer^27,28^.

Our GBO model system maintains tumor-associated macrophages/microglia, endothelial cells, and other features of the tumor microenvironment, such as hypoxia gradients^11^. This provides a unique opportunity to study the interaction between tumor and non-tumor cells and the influence of hypoxia on cell behavior and localization and how this may lead to treatment resistance. Importantly, our procedure to generate organoids directly from micro-dissected tumor pieces with minimal perturbation could be applied to other adult brain tumors, pediatric brain tumors, and possibly tumors from other tissues to better preserve native cell-cell interactions and aspects of the tumor microenvironment, although the conditions may need optimization.

### Comparison with other human glioblastoma model systems

Several model systems, including adherent and suspension cell lines, patient-derived xenografts (PDX), and cerebral organoids have contributed greatly to our understanding of glioblastoma pathobiology, but have their limitations. Traditional *in vitro* monolayer and tumor sphere cultures offer a readily expandable and fairly uniform population of tumor cells for experimentation, but fail to recapitulate the heterogeneity of their parent tumors because the procedures to generate them involve single-cell dissociation and growth in medium containing serum or exogenous bFGF and EGF. These conditions promote clonal selection and expansion over serial passages, which is not favorable to maintain the cellular diversity of parent tumors. These cultures can also require a substantial amount of time to establish and have limited generation fidelity^29,30^. PDX models are thought to better retain the cellular diversity of glioblastomas by providing an *in vivo* environment for tumor growth through direct injection of dissociated tumor cells into the mouse flank or brain. However, PDX models are restricted by variable engraftment efficiency, interactions with animal cells, limited throughput, and long latency in tumor generation, which limit their usefulness for testing personalized therapies^31^. Cerebral organoids, self-organizing 3D cultures that resemble the developing brain^32^, have been genetically altered to develop oncogenic properties or co-cultured with tumor spheres to model tumor cell invasion^33,34^. While cerebral organoids are useful, readily manipulatable models, they represent only the fetal brain and lack important non-neural cell types, such as immune and endothelial cells, and do not recapitulate the cellular diversity or architecture of glioblastoma tumors.

Previously described patient-derived glioblastoma organoids better preserved cellular heterogeneity through direct culturing of tumor pieces in Matrigel and further growth in medium containing bFGF and EGF^35^. These glioblastoma organoids were shown to contain stem cell heterogeneity and recapitulate hypoxia gradients, but the ability of this glioblastoma organoid system to recapitulate features of parent tumors and to serve as a platform for therapeutic testing is unclear. Other organoid models used bio-printing of glioma cells and extracellular matrix on a chip to enable high-throughput drug testing, but the ability of these organoids to recapitulate the cellular heterogeneity and organization of tumors is uncertain^36^. Perhaps the closest model systems to GBOs are glioblastoma explant and slice cultures which avoid tissue dissociation and maintain tumor tissue as chunks or slices with minimal perturbation to preserve the cellular composition and local architecture of tumors^37,38^. These models have been useful for preserving local vasculature and immune cells, studying responses to drug therapies, and visualizing cell division and migration soon after tumor resection but are limited by short-term survival and poor expandability^37,39,40^. Cell death which occurs during initial processing may also complicate analysis. Our GBO methodology addresses the time restriction of explant and slice cultures as GBOs can be cultured for weeks to months after resection and biobanked for future studies. This allows ample time for genetic, transcriptomic, and immunohistochemical analyses that may better inform which chemotherapeutic drugs or CAR-T immunotherapies should be tested. Together, in contrast to other human glioblastoma models, our procedure to generate GBOs preserves the cellular heterogeneity of parent tumors by avoiding cell dissociation and the addition of exogenous bFGF, EGF, and extracellular matrix, which can cause differential cell survival and clonal selection. GBOs are generated in 2-4 weeks with high fidelity, preserve the local cytoarchitecture and cell-cell interactions of resected glioblastoma tissue, and can be expanded and biobanked for future studies.

### Comparison to other tumor organoid methodologies

Our glioblastoma organoid methodology differs fundamentally from other tumor organoid methodologies. Current tumor organoid protocols generally involve single-cell dissociation of epithelial tumor tissue followed by embedding cells into Matrigel to support growth in medium containing many mitogens and growth factors^9,10^. Although this methodology has successfully generated numerous tumor organoid biobanks, dissociating solid tumors into single cells may bias against survival of sensitive cell types and destroys native cell-cell interactions. Likewise, culturing in the presence of exogenous growth factors, small molecules, or serum can contribute to clonal selection and population drift from the original tumor cell composition. The addition of these components also complicates the interpretation of drug treatments because of potential known or unknown interactions between factors in the media and the drugs being tested. The presence of exogenous extracellular matrix that may not accurately recapitulate the extracellular matrix composition of the native tissue can also have profound effects on cell behavior^41^. Using a chemically defined medium facilitates reproducibility for biological studies and clinical applications.

### Expertise needed to implement the protocol

Close coordination with neurosurgeons and pathologists is necessary to implement our protocol since glioblastoma tissue must be obtained from patients and optimal tissue quality is critical for success. Although not required, prior experience with organoid and T cell culture is helpful for implementing our protocol. Experience with tissue processing, immunohistology, and microscopy is helpful for downstream analyses. We are confident that any motivated individual can reproduce this protocol with access to quality resected patient tumor tissue, the proper resources, and careful attention to details.

### Limitations of glioblastoma organoid culture

Although our GBO model provides a timely platform to study the cellular heterogeneity of glioblastoma tumors and patient-specific responses to targeted therapies, it has limitations. The laboratory must be located at or near a major hospital that performs surgical removal of brain tumors. Successful GBO generation depends heavily on close coordination with neurosurgeons to resect viable tumor tissue *en bloc* without cauterization and with pathologists to quickly confirm the diagnosis and limit the time from resection to tissue processing. Differences in tissue quality greatly affect the fidelity of GBO generation as fresh, carefully resected, cell-dense tumor tissue has a greater success rate then heavily cauterized, suctioned, or highly necrotic and cell sparse tissue. Although GBOs recapitulate the heterogeneous population of cells found in parent glioblastoma tumors, populations may inevitably drift with prolonged culture (> 6 months). Changes in population dynamics may or may not accurately recapitulate population changes expected in real tumors. Ideally, GBOs should be biobanked within 2 months to ensure availability of low passage samples for further experimentation.

Typically, we have had more success generating organoids from treatment-naïve primary glioblastomas than recurrent/residual tumors (Fig. 1b). This is most likely due to poorer tissue quality as such tumors usually have increased necrosis and fewer proliferating cells. Similarly, we have had success generating GBOs from tumors containing a wide range of mutations (Fig. 1c), but have had lower success with tumors containing *IDH1* mutations. These tumors are generally less proliferative and arise from lower grade gliomas^42^ which we have had less success propagating.

Since our method forgoes cell dissociation and GBOs are generated directly from micro-dissected tumor pieces, they retain many non-tumor cells, particularly tumor-associated macrophages/microglia and endothelial cells (Fig. 3b-e). While the immune and endothelial cells generally persist for eight weeks and beyond, they inevitably decrease in the percentage of total cells with time because they are not dividing and undergo senescence (Fig. 3c). Our medium is optimized for glioblastoma cell survival and growth and may not be ideal for immune and endothelial cell survival or proliferation. The presence of endothelial cells is particularly variable across GBO samples due to non-uniform occurrence of blood vessels throughout the tumor bulk and variability in the vascularity of different tumors. To address this limitation, GBOs could be co-cultured with immune and/or endothelial cells to reconstitute the microenvironment. This has already been demonstrated for many cerebral organoid models^43-46^, and thus could be feasible in GBOs.

### Limitations of GBO and CAR-T cell co-culture

Our GBO and CAR-T cell co-culture methodology provides a platform to investigate CAR-T cell invasion into solid tumors and killing efficacy, but is not without its limitations. Our system does not recapitulate CAR-T cell infiltration into tumor tissue from blood vessels as would normally occur with peripheral infusion. Our system is also oversimplified because the CAR-T cells have direct access to the tumor cells in culture, whereas they would need to extravasate from blood vessels and migrate to their target cells *in vivo*. Therefore, the ratio of CAR-T cells to tumor cells may not be representative of the number of CAR-T cells that would reach a solid tumor *in vivo*. To better address these limitations, GBOs can be xenografted into rodent brains, and CAR-T cells could be peripherally infused. However, this method is limited by requiring immunodeficient mice to prevent host rejection of the graft. Although GBOs better recapitulate solid tumors, their larger size limits equal loading of fluorescent dyes or indicators to facilitate quantification of cell killing as is routinely done with single cell suspension killing assays. GBOs are less useful for high-throughput screening of many different CAR-T cells since the best way to analyze CAR-T effectiveness is through manual cryo-sectioning, immunohistology, and imaging. GBOs could be dissociated before co-culture with CAR-T cells, but dissociation fails to recapitulate the three-dimensional properties of solid tumors and could cause differential survival of certain cell populations.

### Experimental design Overview

The first part of the following step-by-step procedure describes how to generate glioblastoma organoids (GBOs) from surgically resected patient glioblastoma tumor tissue, passage GBOs for prolonged culture and expansion, and cryopreserve and thaw GBOs for later experimentation (Figs. 1a, 2). The second part describes how to co-culture GBOs with CAR-T cells and assess target specificity and cell killing efficacy by quantification using immunohistology (Figs. 1a, 5).

### Generation of glioblastoma organoids and biobanking

Human glioblastoma tissue obtained from surgical resection is micro-dissected into pieces and cultured in an optimized medium on an orbital shaker to generate glioblastoma organoids as described previously^11^ and in Steps 1-19 of this protocol (Fig. 2a). GBOs are passaged by cutting into smaller pieces as described in Steps 20-26 and cryopreserved and thawed as described in Steps 27-50 of this protocol. GBO generation is verified visually by observing survival and rounding of tumor pieces as they continuously proliferate. Procedures for processing tumor tissue and GBOs for downstream analysis are describe in Box 1. The maintenance of parent tumor cellular heterogeneity, proliferation rates, and hypoxia gradients in GBOs is analyzed by immunohistology as described in Boxes 2 and 3.

### Preparation of CAR-T cells, co-culture with GBOs, and killing assay

Experimental CAR-T cells that recognize an antigen expressed by glioblastoma cells and control CAR-T cells that recognize an antigen not expressed by glioblastoma cells are recovered and expanded from frozen stocks as described in Steps 51-69 and co-cultured with GBOs for 72 hours as described in Steps 70-88 of this protocol. Culture of GBOs in medium containing CAR-T cells on an orbital shaker models CAR-T cell invasion into solid tumors and migration towards antigen-expressing cells. Downstream analyses is described in Steps 89a-d. CAR-T cell target specificity and killing efficacy is assessed by immunohistology for target antigen loss (e.g. EGFRvIII) and cell death (cleaved-caspase-3) in tumor cells. CAR-T cell activation is assessed by immunohistology for T cell proliferation (CD3^+^KI67^+^) and ELISA for cytokines released by activated T cells (IL-2, IFNγ, TNFα).

### Controls

Pieces of parent tumor should always be sampled (Box 1) to assess how well the resulting GBOs recapitulate the cellular heterogeneity, transcriptome, and/or mutations of the parent tumor. Since they are not clonally derived, GBOs have more inherent variability than cell lines, and thus rely heavily on proper controls, adequate sample sizes, and standardized methods to make informative conclusions. To properly interpret immunohistology results, it is imperative to culture, process, perform immunohistology, and image the samples being compared in parallel. Variations in processing should be minimized as they can lead to differences in antibody labeling and a possible misinterpretation of the results. Always confirm antibody specificity using secondary antibody only controls. For all experiments involving immunohistology, we suggest analyzing at least 3-5 tumor pieces and/or GBOs per sample with 3-5 images each, and for transcriptomic and genomic studies we suggest sequencing multiple tumor pieces and/or GBOs together or as triplicates to account for variability between GBOs.

Assessment of CAR-T cell activation and target cell killing is based on the quantification of T cell invasion and proliferation, target antigen loss in tumor cells, and apoptosis of target tumor cells. Two controls are critical to ensure that the results are specific to the experimental CAR-T cells and their target antigen. The first control is using CAR-T cells which recognize an antigen not expressed by glioblastoma cells. For example, include co-culture of GBOs with CD19 CAR-T cells that recognize the B cell lineage^47^. This will test whether the observed cell killing is a result of the antigen recognized by the experimental CAR as opposed to generalized killing in the presence of T cells. The second control is using a GBO sample that does not contain the antigen that the experimental CAR-T cells recognize. For example, if the CAR-T cells recognize EGFRvIII, a mutant variant of EGFR often found in glioblastomas, include co-culture with GBOs that do not contain cells expressing EGFRvIII. This will test the specificity of the experimental CAR for its antigen. If possible, consider a third control that involves co-culture of non-neoplastic neurons, neurospheres, and/or brain organoids with the experimental CAR-T cells to verify that the target antigen is specific to tumor cells and is not expressed by other brain cells. For ELISAs, it is important to include medium sampled from GBOs and CAR-T cells that were not co-cultured with each other to measure baseline cytokine levels.

### Materials

CRITICAL The reagents and equipment listed in this section have been tested to ensure optimal performance for our protocols. Materials may be substituted for equivalents from other suppliers, but we cannot guarantee that they will perform equally.

### Reagents

- 1X RBC lysis buffer (Thermo Fisher Scientific, cat. no. 00433357)
- 16% (vol/vol) formaldehyde, methanol free (Polysciences, cat. no. 18814-10) CRITICAL Ensure that the formaldehyde solution is methanol-free as methanol can have an impact on the quality of fixation, including interference with membrane bound protein staining, cytoskeletal protein staining, and the stability of fluorescent proteins^48^.
- 2-Mercaptoethanol (Thermo Fisher Scientific, cat. no. 21985023)
- 5-ethynyl-2’-deoxyuridine (EdU; Abcam, cat. no. ab146186)
- Antibiotic-antimycotic (Thermo Fisher Scientific, cat. no. 15240062)
- B27 supplement minus vitamin A (Thermo Fisher Scientific, cat. no. 12587010) CRITICAL Use B27 without vitamin A as vitamin A is readily metabolized to retinoic acid which has been shown to decrease the proliferation of glioma stem cells and promote their differentiation^49^.
- Bovine serum albumin (BSA; Sigma-Aldrich, cat. no. B6917)
- Click-iT EdU Alexa Fluor imaging kit (Alexa Fluor 488, Alexa Fluor 555, and Alexa Fluor 647; Thermo Fisher Scientific, cat. nos. C10337, C10338, and C10340)
- Conflikt quaternary disinfectant (Decon Labs, cat. no. 4101)
- DAPI (Sigma-Aldrich, cat. no. 10236276001)
- Dimethyl sulfoxide (DMSO; Sigma-Aldrich, cat. no. D2650)
- Donkey anti-Goat IgG polyclonal secondary antibody (Alexa Fluor 488, Alexa Fluor 555, and Alexa Fluor 647; Thermo Fisher Scientific, cat. nos. A-11055, A-21432, and A-21447, RRIDs: AB_2534102, AB_2535853, and AB_2535864)
- Donkey anti-Mouse IgG polyclonal secondary antibody (Alexa Fluor 488, Alexa Fluor 555, and Alexa Fluor 647; Thermo Fisher Scientific, cat. nos. A-21202, A-31570, and A-31571, RRIDs: AB_141607, AB_2536180, and AB_162542)
- Donkey anti-Rabbit IgG polyclonal secondary antibody (Alexa Fluor 488, Alexa Fluor 555, and Alexa Fluor 647; Thermo Fisher Scientific cat. nos. A-21206, A-31572, and A-31573, RRIDs: AB_2535792, AB_162543, and AB_2536183)
- Donkey serum (Millipore, cat. no. S30)
- Dulbecco’s modified eagle medium/nutrient mixture F-12 (DMEM/F-12; Thermo Fisher Scientific, cat. no. 11320033)
- Dulbecco’s phosphate-buffered saline with calcium and magnesium (DPBS++; Thermo Fisher Scientific, cat. no. 14040133)
- Dulbecco’s phosphate-buffered saline without calcium and magnesium (DPBS; Thermo Fisher Scientific, cat. no. 14190144)
- DuoSet ELISA Ancillary Reagent Kit 1 (R&D Systems, cat. no. DY007)
- Ethyl alcohol, 95% (vol/vol), 190 proof, pure (Pharmco-Aaper, cat. no. 111000190CSGL)
- EZ-PCR mycoplasma test kit (Biological Industries, cat. no. 2070020)
- Gelatin from cold water fish skin (Sigma Aldrich, cat. no. G7041)
- GenePrint 24 system STR profiling kit (Promega, cat. no. B1870)
- GlutaMAX supplement (Thermo Fisher Scientific, cat. no. 35050061)
- Glycine, ultrapure (Thermo Fisher Scientific, cat. no. 15527013)
- Goat anti-Doublecortin polyclonal antibody (Santa Cruz Biotechnology, cat. no. sc-8066, RRID: AB_2088494)
- Goat anti-Nestin polyclonal antibody (Santa Cruz Biotechnology, cat. no. sc-21247, RRID: AB_650014)
- Goat anti-Sox2 polyclonal antibody (Santa Cruz Biotechnology, cat. no. sc-17320, RRID: AB_2286684)
- Hibernate A medium (BrainBits, cat. no. HA)
- Human IFN-gamma DuoSet ELISA (R&D Systems, cat. no. DY285B-05)
- Human IL-2 DuoSet ELISA (R&D Systems, cat. no. DY202-05)
- Human TNF-alpha DuoSet ELISA (R&D Systems, cat. no. DY210-05)
- Human insulin solution (Sigma-Aldrich, cat. no. I9278)
- Human recombinant interleukin 2 (Sigma Aldrich, cat. no. 11011456001)
- Hydrochloric acid (Sigma-Aldrich, cat. no. H1758)
- Hypoxyprobe kit (Hypoxyprobe, cat. no. HP1-100Kit)
- ImmunoCult-XF T Cell Expansion Medium (StemCell Technologies, cat. no. 10981)
- Isopropanol (Sigma-Aldrich, cat. no. 437522)
- MEM non-essential amino acids solution (MEM-NEAAs; Thermo Fisher Scientific, cat. no. 11140050)
- Mouse anti-BLBP monoclonal antibody (Abcam, cat. no. ab131137, RRID: AB_11157091)
- Mouse anti-CD3 monoclonal antibody (BioLegend, cat. no. 344802, RRID: AB_2043995)
- Mouse anti-EGFR monoclonal antibody (Thermo Fisher Scientific, cat. no. MA1-12693, RRID: AB_1074165)
- Mouse anti-GFAP monoclonal antibody (Millipore, cat. no. MAB360, RRID: AB_11212597)
- Mouse anti-Granzyme B monoclonal antibody (R&D Systems, cat. no. MAB2906, RRID: AB_2263752)
- Mouse anti-Ki67 monoclonal antibody (BD Biosciences, cat. no. 550609, RRID: AB_39377)
- Mouse anti-Phospho-Histone H3 monoclonal antibody (Cell Signaling Technology, cat. no. 9706, RRID: AB_331748)
- Mouse anti-VWF monoclonal antibody (Santa Cruz Biotechnology, cat. no. sc-365712, RRID: AB_10842026)
- N2 supplement (Thermo Fisher Scientific, cat. no. 17502048)
- Neural tissue dissociation kit – postnatal neurons (Miltenyi Biotech, cat. no. 130-094-802)
- Neurobasal medium (Thermo Fisher Scientific, cat. no. 21103049)
- Penicillin-streptomycin (Thermo Fisher Scientific, cat. no. 15070063)
- Plasmocin mycoplasma elimination reagent (InvivoGen, cat. no. ant-mpt-1)
- Rabbit anti-CD31 polyclonal antibody (Abcam, cat. no. ab28364, RRID: AB_726362)
- Rabbit anti-Cleaved Caspase-3 polyclonal antibody (Cell Signaling Technology, cat no. 9661 RRID: AB_2341188)
- Rabbit anti-EGF Receptor vIII monoclonal antibody (Cell Signaling Technology, cat. no. 64952, RRID: AB_2773018)
- Rabbit anti-Hopx polyclonal antibody (Santa Cruz Biotechnology, cat. no. sc-30216, RRID: AB_2120833)
- Rabbit anti-Iba1 polyclonal antibody (Wako, cat. no. 019-19741, RRID: AB_839504)
- Rabbit anti-Ki67 polyclonal antibody (Abcam, cat. no. ab15580, RRID: AB_443209)
- Rabbit anti-Olig2 monoclonal antibody (Abcam, cat. no. ab109186, RRID: AB_10861310)
- Rabbit anti-Phospho-Vimentin monoclonal antibody (Cell Signaling Technology, cat. no. 3877, RRID: AB_2216265)
- Rabbit anti-S100B polyclonal antibody (Sigma-Aldrich, cat. no. S2644, RRID: AB_477501)
- Rat anti-CD3 monoclonal antibody (GeneTex, cat. no. GTX11089, RRID: AB_369097)
- Sodium chloride (Sigma-Aldrich, cat. no. 71376)
- Sucrose (Sigma-Aldrich, cat. no. S0389)
- Tissue freezing medium (General Data, cat. no. 1518313)
- Tris base (Sigma-Aldrich, cat. no. 93362)
- Triton X-100 (Sigma-Aldrich, cat. no. T9284)
- TrueBlack lipofuscin autofluorescence quencher (Biotium, cat. no. 23007)
- Trypan blue stain (Thermo Fisher Scientific, cat. no. T10282)
- Tween 20 (Sigma-Aldrich, cat. no. P1379)
- Ultrapure DNase/RNase-free distilled water (Thermo Fisher Scientific, cat. no. 10977015)
- VECTASHIELD Vibrance antifade mounting medium (Vector Laboratories, cat. no. H170010)
- Y-27632 (StemCell Technologies, cat. no. 72304)

### Equipment

- 1.5 mL microcentrifuge tubes (USA Scientific, cat. no. 1615-5500)
- 30 mm conical tube rack (Fisher Scientific, cat. no. 14-809-44)
- Autoclave
- Benchtop minicentrifuge (Benchmark Scientific, cat. no. C1012)
- Benchtop centrifuge (Fisher Scientific, model no. AccuSpin 400)
- Biological safety cabinet (Baker Company, Sterilgard Series)
- Biopsy cryomold (Tissue-Tek, cat. no. 4565)
- Cell counting slides (Thermo Fisher Scientific, cat. no. C10228)
- Charged microscope slides (Fisher Scientific, cat. no. 22-035813)
- CO_2_ incubator (5% CO_2_, 37°C, 90% humidity; Thermo Fisher Scientific, cat. no. 3310)
- Confocal microscope with 10X, 20X, 40X, and 63X objectives (Zeiss, model no. LSM 800)
- Conical tubes (15 and 50 mL; Corning, cat. nos. 05-538-59A and 05-526B)
- CoolCell cell freezing container (Corning, cat. no. 432001)
- Countess II automated cell counter (Thermo Fisher Scientific, cat. no. AMQAX1000)
- Cryogenic vials (Thermo Fisher Scientific, cat. no. 12-567-501)
- Delicate task wipers (Fisher Scientific, cat. no. 34120)
- Disposable filter units with 0.2-µm PES membrane, 500 ml (Thermo Fisher, cat. no. 566-0020)
- Disposable microtome blades (Sakura Finetek USA, cat. no. 4689)
- Dissection microscope (Zeiss, model no. SteREO Discovery.V8)
- Eppendorf variable adjustable-volume pipettes (P1000, P200, and P20; Fisher Scientific, cat. nos. 13-690-032, 13-690-030, and 13-690-027)
- Filter pipette tips (1000 and 200 µL; USA Scientific, cat. nos. 1126-7810 and 1120-8810)
- Fine forceps – curved/serrated (Fine Science Tools, cat. no. 11065-07)
- Fine tip marking pens (Fisher Scientific, cat. no. 13-379-4)
- Glass culture dish (Sigma Aldrich, cat. no. CLS3160102)
- Hydrophobic barrier PAP pen (Vector Laboratories, cat. no. H-4000)
- Inverted microscope (Zeiss, Axiovert 40 CFL model)
- Isopropanol (Sigma Aldrich, cat. no. 437522)
- Label Printer (Fisher Scientific, cat. no. 19-102-824)
- MACS SmartStrainer, 70 μm (Miltenyi Biotech, cat. no. 130-098-462)
- Microtube rack (Fisher Scientific, cat. no. 22-313630)
- Orbital shaker, CO_2_ resistant (Thermo Fisher Scientific, cat. no. 88881102)
- Phosphate-buffered saline, 10X (Thermo Fisher Scientific, cat. no. AM9625)
- Polystyrene serological pipettes (1, 5, 10, and 25 mL; Fisher Scientific, cat. nos. 13-675-15B, 13-675-22, 13-675-20, and 13-675-30)
- Razor blades (Fisher Scientific, cat. no. 12-640)
- Rectangle cover glasses (Fisher Scientific, cat. no. 12-545-F)
- Repair kit for fine forceps (Fine Science Tools, cat. no. 29000-00)
- Research cryostat (Leica, cat. no. CM3050 S)
- Self-laminating Polyester Cryogenic Laboratory Labels (Fisher Scientific, cat. no. 19-102-744)
- Sharpening stone (Fine Science Tools, cat. no. 29008-12)
- Slide-staining chamber (Electron Microscopy Sciences, cat. no. 62010-37)
- Sterilization pouches (Fisher Scientific, cat. no. 01-812-54)
- Syringe filters, 0.2 μm PES (Thermo Fisher Scientific, cat. no. 42225-PS)
- Syringe filters, 0.2 μm nylon (Thermo Fisher Scientific, cat. no. 42225-NN)
- Ultra-low attachment culture plates (6- and 24-well; Corning, cat. nos. CLS3471 and CLS3473)
- Untreated 6-well culture plates (Corning, cat. no. CLS3736)
- Vannas spring scissors – curved/3mm cutting edge (Fine Science Tools, cat. no. 15000-10)
- Weighing dishes (Sigma-Aldrich, cat. no. Z154873)

### Software

- Illustrator CC (Adobe, https://www.adobe.com/products/illustrator.html, RRID:SCR_010279)
- Image J (NIH, https://imagej.nih.gov/ij/, RRID:SCR_003070)
- Imaris (Bitplane, https://imaris.oxinst.com/packages, RRID:SCR_007370)
- Photoshop CC (Adobe, https://www.adobe.com/products/photoshop.html, RRID:SCR_014199)
- Prism (GraphPad, https://www.graphpad.com/scientific-software/prism/, RRID:SCR_002798)
- Zen 2 Blue (Zeiss, https://www.zeiss.com/microscopy/us/products/microscopesoftware/zen.html, RRID:SCR_013672)

### Reagent setup

#### Tumor tissue

The use of human brain tissue and peripheral blood samples was coordinated by the University of Pennsylvania Tumor Tissue/Biospecimen Bank following ethical and technical guidelines on the use of human samples for biomedical research purposes. Patient glioblastoma tissue and peripheral blood samples were collected at the Hospital of the University of Pennsylvania after informed patient consent under a protocol approved by the University of Pennsylvania’s Institutional Review Board. All patient samples were de-identified before processing. Glioblastoma tumor tissue should be obtained directly from *en bloc* surgical resection and placed in glioblastoma dissection medium (Table 1) on ice during transportation and temporary storage. Tissue processing should begin < 2 hours after tissue removal to maximize viability. For best results, prioritize obtaining and processing tumor pieces that are cell-dense and lack substantial surrounding brain tissue or hemorrhage (Fig. 2c).

**Table 1.** Medium components and recipes

| Medium | Component | Recipe |
| --- | --- | --- |
| Glioblastoma dissection medium | Hibernate A | 490 mL |
|  | GlutaMAX supplement (100X) | 5 mL |
|  | Antibiotic-antimycotic (100X) | 5 mL |
| Glioblastoma organoid (GBO) medium | Dulbecco's modified eagle medium/nutrient mixture F-12 (DMEM/F-12) | 235 mL |
|  | Neurobasal medium | 235 mL |
|  | MEM non-essential amino acids solution (100X) | 5 mL |
|  | GlutaMAX supplement (100X) | 5 mL |
|  | N-2 supplement (100X) | 5 mL |
|  | B-27 supplement (50X), minus vitamin A | 10 mL |
|  | Penicillin-streptomycin (100X) | 5 mL |
|  | 2-mercaptoethanol (1000X) | 500 µL |
|  | Human insulin solution | 125 µL |
| Glioblastoma organoid (GBO) freezing medium | Glioblastoma organoid (GBO) medium | 45 mL |
|  | Dimethyl sulfoxide (DMSO) | 5 mL |
|  | 20 mM Y-27632 dihydrochloride | 25 µL |
| T cell medium | ImmunoCult-XF T Cell Expansion Medium | 495 mL |
|  | Penicillin-streptomycin (100X) | 5 mL |

CAUTION Informed patient consent is required for the use of human tissue and all studies must adhere to institutional and governmental ethical and technical guidelines on the use of human samples for biomedical research purposes.

CAUTION When handling multiple patients’ tumors or regional samples of the same tumor simultaneously, pay extra attention to avoid cross-contamination. We recommend using separate dissection tools, culture vessels, and pipette tips for each sample and only reusing tools after proper cleaning and sterilization by autoclave.

CAUTION Glioblastoma organoids should be tested regularly for mycoplasma contamination which can greatly affect the growth and behavior of tumor cells^50^. Testing the original source material is unreliable since mycoplasma can be diluted when tumor tissue is placed in fresh solutions and tumor tissue may become contaminated during dissection and processing. We recommend testing after tumor pieces have been in culture for at least 1 week using a medium sample that has not been recently replaced. Testing must be performed before biobanking and using samples for experimentation. We include a recommended PCR-based testing kit from Biological Industries, which contains detailed instructions for operation (additional equipment may be required), in the reagents section. Mycoplasma testing can also be outsourced to vendors, including ATCC (atcc.org/services/testing_services/mycoplasma_testing) and Cell Line Genetics (clgenetics.com/sample_mycoplasma_testing) by following the instructions on their websites. GBO samples that test positive for mycoplasma are discarded unless they are particularly important for later clinical testing, for which we attempt mycoplasma removal by treatment with Plasmocin (InvivoGen) using the instructions provided with the reagent.

CAUTION To verify that the glioblastoma tissue and resulting organoids are matched with the correct patient medical information and that the samples are not cross-contaminated we recommend routine short-tandem repeat (STR) profiling of DNA isolated from patients and GBOs. This is especially important as proper quality control when creating a biobank. We include a recommended STR profiling kit from Promega (additional equipment may be required), which contains detailed instructions for operation, in the reagents section. STR profiling can also be outsourced to vendors, including ATCC (atcc.org/services/testing_services/cell_authentication_testing_service) and Cell Line Genetics (clgenetics.com/our-services/short-tandem-repeat-analysis) by following the instructions on their websites.

### Antibody dilution solution

Add 100 μL of Triton X-100 and 5 mL of donkey serum to 95 mL of TBST and mix thoroughly by inverting or using a magnetic stir bar and plate. Sterilize by syringe or vacuum filtration through a 0.2 μm PES membrane. Prepare single-use aliquots and store at ≤ −20°C for up to 6 months. Avoid repeated freeze-thaw cycles.

### Autofluorescence quenching solution

Dilute 20X stock TrueBlack reagent with 70% (vol/vol) ethanol to obtain a 1X solution. Prepare the volume needed immediately before use and protect from exposure to light.

CAUTION Ethanol is highly flammable and can cause irritation when inhaled; keep away from flames and use in well-ventilated areas.

### Blocking and permeabilization solution

Dissolve 0.1 g of gelatin, 2.25 g of glycine, and 1 g of bovine serum albumin (BSA) in 90 mL of TBST. Add 0.5 mL Triton X-100 and 10 mL donkey serum and mix the solution well by inverting or using a magnetic stir bar and plate. Sterilize by syringe or vacuum filtration through a 0.2 μm PES membrane. Prepare single-use aliquots and store at ≤ −20°C for up to 6 months. Avoid repeated freeze-thaw cycles.

### EdU solution

Dissolve 50 mg of EdU in 19.8 mL of DMSO to obtain a 10 mM stock solution. Sterilize by syringe or vacuum filtration through a 0.2 μm nylon membrane. Prepare single-use aliquots and store at ≤ −20°C for up to 6 months. Avoid repeated freeze-thaw cycles.

CAUTION EdU is a chemical agent which incorporates into newly synthesized DNA. Use appropriate personal protective equipment to avoid direct contact with EdU solutions.

### Ethanol mixture (70% vol/vol)

Add 737 mL of 190 proof ethanol to a 1 L graduated cylinder. Fill the graduated cylinder to 1 L with ultra-pure water. Store at room temperature in a sealed container or spray bottle.

CAUTION Ethanol is highly flammable and can cause irritation when inhaled; keep away from flames and use in well-ventilated areas.

### Formaldehyde solution (4% vol/vol)

Dilute 10 mL of 16% methanol-free formaldehyde solution to 8% by adding 10 mL of 2X TBS. Dilute the 8% solution to 4% by adding 20 mL of 1X TBS. Store at 4°C for up to 1 week.

CAUTION Formaldehyde is an irritant and known carcinogen and should only be handled with proper personal protective equipment within a well-ventilated area.

### Glioblastoma dissection medium

Combine 490 mL Hibernate A, 5 mL GlutaMAX supplement (100X), and 5 mL antibiotic-antimycotic (100X) (Table 1). Sterilize by vacuum filtration through a 0.2 μm PES membrane. Store at 4°C for up to 4 weeks.

CRITICAL Antibiotic-antimycotic is important as prophylaxis for bacterial or fungal contamination during tissue processing by pathology and transportation to the laboratory.

### Glioblastoma organoid medium

Combine 235 mL DMEM:F12 medium, 235 mL Neurobasal medium, 5 mL MEM non-essential amino acids solution (100X), 5 mL GlutaMAX supplement (100X), 5 mL penicillin-streptomycin (100X), 5 mL N-2 supplement (100X), 10 mL B-27 minus vitamin A supplement (50X), and 125 μL human recombinant insulin (Table 1). Sterilize by vacuum filtration through a 0.2 μm PES membrane. Store at 4°C for up to 4 weeks. Add the needed amount of 2-mercaptoethanol (1000X) to an aliquot of medium immediately before changing GBO medium.

CAUTION 2-mercaptoethanol is not stable in solution. Add fresh, sterile 2-mercaptoethanol to an aliquot of medium before each medium change. Store 1000X 2-mercaptoethanol solution at 4°C until date indicated on bottle.

### Glioblastoma organoid freezing medium

Add 5 mL dimethyl sulfoxide (DMSO) to 45 mL glioblastoma organoid medium (Table 1). Sterilize by syringe or vacuum filtration through a 0.2 μm PES membrane. Store at 4°C for up to 2 weeks.

### Isopropanol cooled with dry ice

Add dry ice to a suitable ice bucket. Slowly add isopropanol to completely submerge the dry ice pieces. Allow isopropanol to cool for 5 minutes before using. Use isopropanol cooled with dry ice to flash freeze samples within 15 minutes.

CAUTION Isopropanol is highly flammable and can cause irritation when inhaled; keep away from flames and use in well-ventilated areas.

CAUTION Dry ice and isopropanol cooled with dry ice are very cold. Use appropriate personal protective equipment to avoid injury.

CAUTION Isopropanol may remove ink labels on tubes or cryomolds. Use an isopropanol-resistant marker to prevent loss of sample identity.

### Pimonidazole-HCl solution

Dissolve 100 mg of pimonidazole-HCl in 860 μL of DPBS to obtain a 400 mM stock solution. Sterilize by syringe or vacuum filtration through a 0.2 μm PES membrane. Prepare single-use aliquots and store at ≤ −20°C for up to 6 months. Avoid repeated freeze-thaw cycles and protect from prolonged exposure to light.

### Sucrose solution (30% wt/vol)

Dissolve 30 g of sucrose in 90 mL of TBS. Adjust final volume to 100 mL with TBS and sterilize by syringe or vacuum filtration through a 0.2 μm PES membrane. Store at 4°C for up to 6 months.

### T cell medium

Add 5 mL of penicillin-streptomycin solution to 495 mL of ImmunoCult-XF T cell expansion medium (Table 1). Sterilize by syringe or vacuum filtration through a 0.2 μm PES membrane. Store at 4°C for up to 4 weeks.

### Tris-buffered saline (10X)

Dissolve 24 g of tris base and 88 g of sodium chloride in 900 mL of ultra-pure water. Adjust pH to 7.6 with 12 N hydrochloric acid. Adjust final volume to 1 L with ultra-pure water and sterilize by autoclaving. Store stock solution at 4°C for up to 3 months. Dilute stock solution in ultra-pure water to obtain a 1X working solution. Store working solution at room temperature for up to 1 month.

### Tris-buffered saline Tween-20 (10X)

Dissolve 24 g of tris base, 88 g of sodium chloride, and 10 mL of Tween-20 in 900 mL of ultra-pure water. Adjust pH to 7.6 with 12 N hydrochloric acid. Adjust final volume to 1 L with ultra-pure water and sterilize by autoclaving. Store stock solution at 4°C for up to 3 months. Dilute stock solution in ultra-pure water to obtain a 1X working solution. Store working solution at room temperature for up to 1 month.

### Wide bore P1000 pipette tips

Carefully cut the ends of P1000 pipette tips using a sterile razor blade or scissors to create an approximately 3 mm diameter opening. Place pipette tips into a pipette tip box and autoclave.

### Y-27632 dihydrochloride

Dissolve 50 mg Y-27632 in 7.8 mL of DMSO to obtain a 20 mM stock solution. Sterilize by syringe filtration through a 0.2 μm nylon membrane. Store at ≤ −20°C for up to 12 months. Avoid repeated freeze-thaw cycles and protect from prolonged exposure to light.

## Procedures

### Establishment of glioblastoma organoid culture from resected patient tumor tissue (Timing 2-4 h for tissue processing, 2-4 w for organoid establishment)

CRITICAL Optimal tumor tissue is resected *en bloc* from cell-dense regions without substantial necrosis or surrounding brain tissue. Cell-dense tumor tissue is identified as solid white in appearance with occasional blood vessels. Contaminating brain tissue may be difficult to identify but usually appears as pink-yellow tissue that is less cell-dense and may have viable tumor tissue attached. Tumor tissue with substantial hemorrhage is easily identified as pieces with dark red areas of clotted blood. Necrotic tissue is loosely organized, darker, and falls apart easily during dissection. Example images of cell-dense tumor tissue, contaminating brain tissue, and tumor tissue with substantial hemorrhage are provided in Fig. 2c. Tumor tissue must be processed immediately after resection and confirmed pathologic diagnosis to maximize cell viability. The major steps and conditions for processing resected glioblastoma tissue into tumor pieces for GBO culture are outlined in Fig. 2a.

CRITICAL Perform all steps within a sterile laminar flow biosafety cabinet to minimize risk of microbial contamination. Sterilize all dissection tools and pipette tips by autoclave. Ensure that all non-autoclavable items and equipment within the biosafety cabinet are disinfected with 70% ethanol before proceeding.

1. After preliminary diagnosis of glioblastoma by an attending neuropathologist, transfer surgically resected glioblastoma tumor tissue to a 15 mL conical tube containing 5 mL ice cold glioblastoma dissection medium (Table 1) for transportation and short-term storage (Fig. 2b). CRITICAL STEP Obtain proper institutional review board approval and informed patient consent before the surgery and follow all ethical guidelines for the use of human tissue. CRITICAL STEP Begin processing the tumor tissue within 2 hours after initial resection to maximize tissue viability. Leaving the tissue overnight at 4°C will substantially decrease tissue viability. CRITICAL STEP If the surgical resection is large, the tumor tissue can be subdivided into regional samples to analyze intra-tumoral heterogeneity. Subdivide the tumor tissue in a laminar flow biosafety cabinet with a sterile razor blade or scalpel, and place regional samples into separate 15 mL conical tubes with 5 mL ice cold glioblastoma dissection medium.
2. Place conical tubes with tumor pieces into a tube rack and carefully aspirate medium above sunken tumor pieces.
3. Wash the tumor tissue three times with 10 mL ice cold DPBS++ by letting tumor pieces sink to the bottom of the conical tube before carefully aspirating the supernatant (Fig. 2b). CRITICAL STEP Direct exposure to blood components is harmful to cells in culture. Thorough washing is critical to preserving the viability of tumor cells.
4. Transfer tumor pieces to a sterile glass dissection dish containing 10 mL ice cold glioblastoma dissection medium within a sterile laminar flow biosafety cabinet with a suitable dissection microscope (Fig. 2c). CRITICAL STEP Use glass dishes to avoid introducing debris from scratched plastic dishes into the cultures during dissection. CRITICAL STEP Glass dishes may be reused after washing and autoclaving. Rinse glass dishes thoroughly with deionized water after washing and before autoclaving to remove residual detergents which are toxic to living cells.
5. Carefully cut the tumor tissue into 0.5 – 1 mm pieces using fine spring dissection scissors and forceps (Movie 1). Start by cutting the entire tissue into large pieces. This quickly exposes the tissue to nutrients within the medium. Then cut each large piece into smaller pieces (Fig. 2d). CRITICAL STEP Do not shred the tissue by pulling it apart. Only make straight cuts using sharpened dissection scissors. Dissection scissors and forceps can be sharpened manually using the sharpening stone and repair kit listed in the materials section or by sending to Corte Instruments (corteinstruments.com). TROUBLESHOOTING
6. While dissecting the tissue, remove pieces with substantial necrosis, hemorrhage, or brain tissue as these pieces have sparse amounts of viable tumor cells and do not produce satisfactory organoids. CRITICAL STEP Pieces with substantial necrosis are identified as darker with loose organization that falls apart easily during dissection. Pieces with substantial hemorrhage are identified as dark red and consisting mostly of clotted blood (Fig. 2c). Brain tissue usually appears as pink-yellow tissue that is less cell-dense (Fig. 2c). Brain tissue may be difficult to distinguish from tumor tissue at first but will be easier to identify as pieces that fail to grow and round up after 1-2 weeks (Fig. 2f, i).
7. Collect micro-dissected tumor pieces into a 15 mL conical tube using a P1000 pipette with a wide bore tip and carefully aspirate the medium above sunken tumor pieces.
8. Wash the tumor pieces three times with 10 mL room temperature DPBS++ by letting the tumor pieces sink to the bottom of the conical tube before carefully aspirating the supernatant.
9. Collect and process a sample of tumor pieces for downstream analysis as described in Box 1. CRITICAL STEP Sampling micro-dissected tumor pieces is important for comparing the cellular heterogeneity, transcriptional signatures, and mutation profiles to the resulting GBOs. The micro-dissected tumor pieces are a more accurate reference than tissue banked by pathology since GBOs are generated from the same tumor sample.
10. Add 10 mL of red cell lysis buffer and incubate the conical tube on a rocking platform shaker set to approximately 10 tilt cycles per minute at room temperature for 10 minutes to lyse the majority of contaminating red blood cells. The buffer may turn red if the tumor sample contains substantial blood.
11. Carefully aspirate red cell lysis buffer and wash the tumor pieces three times with 10 mL room temperature DMEM:F12 by letting tumor pieces sink to the bottom of the conical tube before carefully aspirating the supernatant.
12. Transfer the dissected tumor pieces to a 6-well flat-bottom, non-treated culture plate with 10-20 tumor pieces and 4 mL of GBO medium (Table 1) per well.
13. Culture the tumor pieces on a CO_2_ resistant orbital shaker rotating at 120 rpm within a 37°C, 5% CO_2_, 90% relative humidity sterile incubator.
14. Replace approximately 75% of the medium every 48 hours by tilting the culture plate at a 45° angle and aspirating the medium above the sunken tumor pieces/GBOs.
15. After 1 week of culture, many pieces should appear to round up (Fig. 2e). Remove tumor pieces that do not round up and are cell-sparse (Fig. 2f). These pieces are often composed of cell-sparse extracellular matrix from resected brain or necrotic tissue and will not form GBOs. CRITICAL STEP Within the first week of culture the tumor pieces will often shed cellular and blood debris making the medium cloudy. The shedding should soon cease as the tumor pieces form rounded organoids generally within 2 weeks depending on tissue quality and patient-specific tumor growth characteristics.
16. Continue culturing GBOs for 2-4 weeks. By this point it should be readily apparent which pieces are growing by their round, cell-dense morphology (Fig. 2g). Continue to remove pieces that do not grow until the cultures consist entirely of round GBOs. CRITICAL STEP The criteria for successful establishment of GBOs from a given patient’s tumor is that the micro-dissected tumor pieces survive for 2 weeks, develop a spherical morphology, and continue growing in culture (Fig. 2g). The quality of GBOs and the extent to which they recapitulate the parent tumor are assessed by immunohistology (Box 2, Fig. 3a) showing maintenance of proliferation and cell markers and transcriptional and genomic analyses showing maintenance of gene expression and mutations (refer to Jacob et al., 2020)^11^. CRITICAL STEP Cell-dense GBOs appear lighter by bright-field microscopy than necrotic tissue, although given enough time in culture, some cell-sparse necrotic pieces may become surrounded by growing tumor cells (Fig. 2i). TROUBLESHOOTING
17. Sample GBOs at different timepoints (e.g. 1 week, 2 weeks, etc.) and process for downstream analysis as described in Box 1, and immunohistology as described in Box 2.
18. (Optional) Analyze hypoxia gradients and active tumor cell proliferation by incubation of live GBOs with pimonidazole-HCl and EdU respectively as described in Box 3.
19. (Optional) Monitor GBO growth by bright-field imaging of individual GBOs every week and tracking their relative size over time (Fig. 1e).

### Passaging glioblastoma organoids (Timing 1-2 h)

CRITICAL GBOs should be passaged once they reach approximately 1-2 mm in diameter (Fig. 2h) to avoid substantial necrotic cell death within the core.

CRITICAL Dissection tools must be thoroughly washed and sterilized by autoclave between handling of different samples to avoid microbial contamination and cross-contamination between samples. Dissection scissors should be sharpened routinely to facilitate cutting without shredding the tissue.

20. Transfer GBOs to a 15 mL conical tube using a P1000 pipette with a wide bore tip.
21. Aspirate medium and wash once with glioblastoma dissection medium by letting GBOs sink to the bottom of the conical tube before carefully aspirating the supernatant.
22. Transfer GBOs to a glass dissection dish containing 10 mL glioblastoma dissection medium within a laminar flow biosafety cabinet with a suitable dissection microscope.
23. Dissect each GBO into four pieces by sequential bisection using fine spring dissection scissors (Fig. 2k, l). CRITICAL STEP Passaging by sequential bisection avoids cell dissociation that can lead to clonal selection and preserves the cellular heterogeneity by maintaining cells from the core and periphery of the GBO in each piece. Passaging using enzymatic or non-enzymatic dissociation will destroy cell-cell interactions within GBOs. TROUBLESHOOTING
24. Collect GBO pieces in a 15 mL conical tube and wash three times with 10 mL DMEM:F12 by letting GBO pieces sink to the bottom of the conical tube before carefully aspirating the supernatant.
25. Transfer the GBO pieces to a 6-well flat-bottom, non-treated culture plate with 10-20 GBO pieces and 4 mL of GBO medium per well.
26. Return GBO pieces to culture on a CO_2_ resistant orbital shaker rotating at 120 rpm within a 37°C, 5% CO_2_, 90% relative humidity sterile incubator.

### Cryopreservation of glioblastoma organoids (Timing 2-3 h)

CRITICAL GBOs should be cryopreserved within 4-8 weeks after successful generation to preserve low passage samples for later experimentation.

CRITICAL Each GBO sample should undergo short tandem repeat (STR) profiling and mycoplasma testing before cryopreservation (recommendations for mycoplasma testing and STR profiling are described in the reagent setup section for tumor tissue). Mycoplasma can greatly affect tumor cell behavior^50^ and any GBO samples that test positive for mycoplasma are unusable for experimentation and should be disposed of or quarantined to prevent cross-contamination to other samples.

27. Transfer GBOs to a 15 mL conical tube using a P1000 pipette with a wide bore tip.
28. Aspirate medium and wash once with glioblastoma dissection medium by letting GBOs sink to the bottom of the conical tube before carefully aspirating the supernatant.
29. Transfer GBOs to a glass dissection dish containing 10 mL glioblastoma dissection medium within a laminar flow biosafety cabinet with a suitable dissection microscope.
30. Dissect GBOs into approximately 0.5 mm diameter pieces by sequential bisection using fine spring dissection scissors. CRITICAL STEP GBOs are cut to pieces no larger than 0.5 mm in diameter to facilitate perfusion with GBO freezing medium (Table 1). The size of GBO pieces is critical for cryopreservation as larger pieces do not recover well.
31. Collect GBO pieces in a 15 mL conical tube and wash three times with 10 mL DMEM:F12 by letting GBO pieces sink to the bottom of the conical tube before carefully aspirating the supernatant.
32. Transfer the GBO pieces to a 6-well flat-bottom, non-treated culture plate with 10-20 GBO pieces and 4 mL of GBO medium with 10 μM Y-27632 per well.
33. Culture the tumor pieces on a CO_2_ resistant orbital shaker rotating at 120 rpm within a 37°C, 5% CO_2_, 90% relative humidity sterile incubator for 60 minutes. CRITICAL STEP Incubation with 10 μM Y-27632 helps to increase cell viability during freezing.
34. Collect GBO pieces in a 15 mL conical tube using a P1000 pipette with a wide bore tip.
35. Aspirate GBO medium with 10 μM Y-27632, resuspend GBO pieces in 10 mL GBO freezing medium (Table 1), and incubate on a rocking platform shaker set to approximately 10 tilt cycles per minute at room temperature for 5 minutes to let GBO freezing medium fully perfuse the organoids. CRITICAL STEP In contrast to single-cell suspensions, GBO pieces are large and require a short incubation step in freezing medium to allow DMSO-containing medium to better penetrate cells. This incubation step helps to increase cell viability during freezing.
36. Aspirate GBO freezing medium and resuspend GBO pieces in 2 mL fresh GBO freezing medium.
37. Transfer approximately 20 GBOs into each cryogenic vial containing 1 mL of GBO freezing medium. CRITICAL STEP Pre-label cryogenic vials with sample identifying information, age, passage number, and date. We recommend temperature and ethanol resistant labels to avoid loss of information during liquid nitrogen storage and vial recovery.
38. Place cryogenic vials containing GBO pieces into a CoolCell freezing container (any cell freezing container that cools at a rate of −1°C per minute is suitable) and place the container into a −80°C freezer overnight. CRITICAL STEP Do not snap-freeze cryovials as this will lead to poor viability.
39. Transfer the cryogenic vials containing GBO pieces to the air phase of liquid nitrogen (≤ −135°C) for long-term storage. CRITICAL STEP Long-term storage in a temperature-stable liquid nitrogen tank is essential for frozen tissue viability. Avoid large fluctuations in temperature and always have a back-up plan for when the liquid nitrogen source or tank malfunctions. PAUSE POINT Cryopreserved GBOs can be stored in liquid nitrogen indefinitely. We have successfully recovered GBOs cryopreserved for over 2 years.

### Thawing glioblastoma organoids (Timing 1 h)

40. Locate cryogenic vials from liquid nitrogen storage and place them on dry ice.
41. Immediately thaw vials in a 37°C water bath, gently swirling the tubes to ensure even thawing. CRITICAL STEP Vials should be thawed quickly and evenly at 37°C and not left to warm past thawing to increase the viability of recovered GBOs.
42. Remove vials from the 37°C water bath when only a small chunk of ice remains.
43. Sterilize vials with 70% ethanol and place in a tube rack within a sterile biosafety cabinet.
44. Gently transfer GBOs to a 50 mL conical tube using a P1000 pipette with a wide bore tip.
45. Add 10 mL of GBO medium with 10 μM Y-27632 dropwise while swirling the tube to slowly dilute the DMSO-containing GBO freezing medium. CRITICAL STEP Gradual recovery by slowly diluting away the DMSO within the freezing medium improves the viability of thawed organoids.
46. Aspirate the supernatant and add 2 mL of GBO medium with 10 μM Y-27632.
47. Transfer the GBO pieces to a 6-well flat-bottom, non-treated culture plate with 10-20 GBO pieces and 4 mL of GBO medium with 10 μM Y-27632 per well (Fig. 2m; left).
48. Culture GBO pieces overnight in a 37°C, 5% CO_2_, 90% relative humidity sterile incubator without shaking.
49. The next day, replace medium with GBO medium without 10 μM Y-27632 and transfer the culture plate to a CO_2_ resistant orbital shaker rotating at 120 rpm within a 37°C, 5% CO_2_, 90% relative humidity sterile incubator.
50. Allow GBOs to recover for 1-2 weeks before using for experimentation. Properly cryopreserved and thawed GBOs should have a round morphology after 1-2 weeks (Fig. 2m; middle) and continue growing for over 4 weeks (Fig. 2m; right). TROUBLESHOOTING

### Preparation of CAR-T cells (Timing 1 h for thawing, 3 days for recovery and expansion)

CRITICAL All CAR-T cells being compared must be prepared in parallel to limit variations in processing that could lead to inaccurate interpretations.

CRITICAL The CAR-T cells used to generate the sample data in Fig. 5 were generated by transducing T cells isolated from donor leukapheresis products with genome integrating lentiviruses harboring the CAR sequence. The Supplemental Methods file includes information on the materials and procedures used to generate the CAR-T cells. Not included in this protocol are detailed procedures for generating CAR-T cells as this is beyond our scope and already described in the literature^22,51^.

CRITICAL T cell batches with roughly 30% expressing the CAR are used in our experiments. Keep this is mind when interpreting CAR-T cell to tumor cell ratios.

51. Prepare culture vessel by adding 10 mL of T cell medium (Table 1) with 100 units/mL interleukin-2 (IL-2) to a T75 flask and leaving the T75 flask in a 37°C, 5% CO_2_, 90% relative humidity sterile incubator to equilibrate until Step 61. Prepare a separate T75 flask for each different CAR-T cell you will use for co-culture with GBOs. CRITICAL STEP Prepare T cell medium with fresh IL-2 by adding single-use sterile aliquots of IL-2 stored at ≤ −80°C to the appropriate amount of T cell medium and store at 4°C for ≤ 1 week to ensure optimal cytokine activity. Avoid pre-warming media to 37°C to minimize cytokine degradation. Consistent storage and usage of IL-2 is essential to obtaining optimal and reproducible results.
52. Locate cryogenic vials containing appropriate control and target CAR-T cells from liquid nitrogen storage and place them on dry ice. Refer to the Controls section in Experimental Design for suggested controls to include with each experiment.
53. Immediately thaw vials in a 37°C water bath, gently swirling the tubes to ensure even thawing. CRITICAL STEP Vials should be thawed quickly and evenly at 37°C and not left to warm past thawing to increase the viability of recovered cells.
54. Remove vials from the 37°C water bath when only a small chunk of ice remains.
55. Sterilize vials with 70% ethanol and place in a tube rack within a sterile biosafety cabinet.
56. Gently transfer contents of cryogenic vials to a 50 mL conical tube using a P1000 pipette. CRITICAL STEP Use separate pipette tips, conical tubes, serological pipettes, and aspiration pipettes for each different type of CAR-T cell to avoid cross-contamination between samples.
57. Add 10 mL of T cell medium dropwise while swirling the tube to slowly dilute the DMSO-containing freezing medium. CRITICAL STEP Gradual recovery by slowly diluting away the DMSO within the freezing medium improves the viability of thawed CAR-T cells.
58. Centrifuge conical tubes containing CAR-T cells at 300 g for 5 minutes to pellet.
59. After centrifugation, carefully aspirate the supernatant and add 5 mL of T cell medium with 100 units/mL IL-2 to the conical tube.
60. Gently resuspend the CAR-T cell pellet using a 5 mL pipette and pipette aid.
61. Transfer 5 mL of CAR-T cells to the T75 flasks prepared in step 51.
62. Culture flasks for 24 hours in a 37°C, 5% CO_2_, 90% relative humidity sterile incubator. CRITICAL STEP Culturing the thawed CAR-T cells for 24 hours in T cell medium with 100 units/mL IL-2 greatly increases cell viability and expansion. PAUSE POINT CAR-T cells are cultured in T cell medium with 100 units/mL IL-2 for 24 hours before proceeding.
63. Prepare new T75 flasks with 10 mL of T cell medium without IL-2 for each CAR-T cell flask cultured in Step 62 and leave the T75 flasks in a 37°C, 5% CO_2_, 90% relative humidity sterile incubator to equilibrate until Step 68.
64. Collect CAR-T cells from T75 flasks into 15 mL conical tubes.
65. Centrifuge conical tubes containing CAR-T cells at 300 g for 5 minutes to pellet.
66. After centrifugation, carefully aspirate the supernatant and add 5 mL of fresh T cell medium without IL-2 to each conical tube.
67. Gently resuspend the CAR-T cell pellets using a 5 mL pipette and pipette aid.
68. Transfer 5 mL of CAR-T cells to the T75 flasks prepared in Step 63.
69. Culture flasks for an additional 48 hours in a 37°C, 5% CO_2_, 90% relative humidity sterile incubator. CRITICAL STEP Culturing the CAR-T cells for 48 hours in T cell medium without IL-2 is necessary to reduce the effects of IL-2 stimulation. This is imperative when comparing CAR-T cell proliferation and cytokine release between control and target antigen groups as IL-2 is a potent CAR-T cell activator that is released when CAR-T cells engage their target antigen. CRITICAL STEP After 48 hours sample 1 mL of the CAR-T cell medium for mycoplasma testing. Verify that each batch of CAR-T cells is negative for mycoplasma before proceeding. Do not sample before 48 hours as mycoplasma may be difficult to detect soon after thawing. PAUSE POINT CAR-T cells are cultured in T cell medium for 48 hours before proceeding. TROUBLESHOOTING

### Co-culture of GBOs with CAR-T cells (Timing 1-5 days)

CRITICAL Verify expression of the target antigen in GBOs by immunohistology as described in Box 2 before proceeding with co-culture (Fig. 5b).

70. Transfer GBOs to a 15 mL conical tube and carefully aspirate medium. Use a separate conical tube for different GBO samples.
71. Wash GBOs three times with 10 mL DMEM:F12 by letting GBOs sink to the bottom of the conical tube before carefully aspirating the supernatant.
72. Add 5 mL of T cell medium to the 15 mL conical tube. CRITICAL STEP The CAR-T cell and GBO co-culture assay is performed in T cell medium as reported previously for cytotoxicity assays in the literature^22^ to ensure optimal activity of the T cells. GBOs tolerate T cell medium during the duration of this experiment with no obvious increase in cell death in control conditions (Fig. 5h). GBO proliferation may decrease due to the absence of insulin and other nutrients, but this does not affect the experimental purpose of assessing CAR-T cell target specificity and killing.
73. Transfer a single GBO into each well of a 24-well ultra-low attachment culture plate containing 250 μL of T cell medium using a P1000 pipette with a wide bore tip. Use at least 3 GBOs for each treatment group but keep individual GBOs in separate wells. CRITICAL STEP Use similarly sized GBOs between samples and treatment groups to minimize the effects of initial size on the experiment outcomes. Because measuring the size of GBOs by volume or weight is difficult, estimate the size of GBOs by taking a bright-field image and measuring the 2D area. CRITICAL STEP Place each GBO into an individual well to control and account for differences between GBOs.
74. Place 24-well culture plates containing GBOs onto a CO_2_ resistant orbital shaker rotating at 120 rpm within a 37°C, 5% CO_2_, 90% relative humidity sterile incubator until Step 83.
75. Separately dissociate three similarly sized GBOs by following the instructions included with the postnatal neuron dissociation kit (additional materials may be required). CRITICAL STEP Enzyme incubation timing may need to be optimized for each sample to reach a single-cell suspension.
76. Perform a cell count manually or by following the instructions included with an automated cell counter to estimate the cell number in each GBO (Fig. 1f). CRITICAL STEP The cell count within each similarly sized GBO is important when determining how many CAR-T cells to add to reach the desired CAR-T cell to tumor cell ratio.
77. Collect CAR-T cells from T75 flasks into 15 mL conical tubes.
78. Centrifuge conical tubes containing CAR-T cells at 300 g for 5 minutes to pellet.
79. After centrifugation, carefully aspirate the supernatant and add 2 mL of fresh T cell medium to each conical tube.
80. Gently resuspend the CAR-T cell pellets using a P1000 pipette.
81. Count the cell number manually or by following the instructions included with an automated cell counter. CRITICAL STEP Include Trypan Blue stain when counting cells to assess the percentage of dead cells. A count of >5-10% dead cells is a sign of unhealthy T cells that may lead to suboptimal results.
82. Dilute CAR-T cell suspension so that 250 μL contains 1/10 the number of CAR-T cells as the estimated cell count in each GBO obtained in Step 76 to achieve a 1:10 CAR-T cell to GBO cell ratio. CRITICAL STEP The ratio of CAR-T cells to GBOs can be adjusted, but keep in mind that a higher ratio will likely result in more rapid killing. The optimal ratio may need to be determined by experimentation.
83. Add 250 μL of diluted CAR-T cell suspension to each well of 24-well plates containing GBOs prepared in Steps 70-74.
84. Transfer the culture plate to a CO_2_ resistant orbital shaker rotating at 120 rpm within a 37°C, 5% CO_2_, 90% relative humidity sterile incubator.
85. Co-culture GBOs with CAR-T cells for 1-5 days and process for immunohistology as described in Box 1. GBOs co-cultured with CAR-T cells may be sampled at multiple timepoints to access the time course of CAR-T cell invasion and killing. TROUBLESHOOTING
86. (Optional) Take bright-field images of GBOs at sequential time points to observe gross changes in T cell number and GBO health (Fig. 5c, d).
87. (Optional) Sample 50 μL of medium from each well for cytokine measurements by ELISA at different time points (Fig. 5e) (See Downstream Assays, Step 89a). We recommend collecting medium samples at least every 24 hours to obtain temporal measurements. Replace the medium removed from each well with 50 μL of fresh T cell medium.
88. (Optional) Perform immunohistology on GBOs as described in Box 2 and quantify T cell proliferation, tumor cell death, and target antigen loss (Fig. 5f-h) (See Downstream Assays, Steps 89b-c).

### Downstream assays (Timing variable)

89. During co-culture, medium can be sampled and evaluated for cytokines released during T cell activation (option a). After co-culture, samples can be processed for downstream analysis (Box 1) and analyzed by immunohistology (Box 2) to assay for T cell invasion and proliferation (option b), target tumor antigen loss (option c), and tumor cell killing (option d). CRITICAL STEP All samples that are being compared should be processed in parallel using identical conditions to limit the effect of differences in preparation on result interpretation.
  a. **Analysis of cytokines released by activated T cells**
    i. Aliquot 50 μL of T cell medium and GBO medium separately as negative controls. CRITICAL STEP Always include negative controls with every experiment to control for any cytokines already present in the media.
    ii. Collect 50 μL of medium from each well containing individual GBOs in T cell medium from Step 73 as the baseline sample in a 1.5 mL microcentrifuge tube.
    iii. Collect 50 μL of medium after the addition of CAR-T cells and after every 12-24 hours.
    iv. Add 50 μL of T cell medium to each well to make up for the volume sampled. CRITICAL STEP Replacing sampled medium with fresh medium is critical to avoid incidental concentration of cytokines.
    v. Centrifuge 1.5 mL microcentrifuge tubes at 2000 g for 5 minutes to pellet cells and debris. CRITICAL STEP Removing cell debris improves reliability of ELISA reaction and ensures that only extracellular cytokines are measured.
    vi. After centrifugation, carefully aspirate the supernatant and add to a new 1.5 mL centrifuge tube.
    vii. Snap-freeze the tubes in dry ice-isopropanol and store in a −80°C freezer until ready for analysis. PAUSE POINT Frozen medium samples can be stored at −80°C for over 2 months. Alternatively, fresh medium samples can be stored at 4°C for up to three days before analysis.
    viii. Thaw medium samples on ice and perform ELISA on 10-fold diluted medium using the instructions included with the ELISA kit (Fig. 5e). Recommended ELISA kits are included in the Materials section (additional materials may be required), although other ELISA kits may be used as a substitute. CRITICAL STEP In most cases it is necessary to dilute the medium 10-fold to remain within the working detection range of the ELISA. Optimization using serial dilutions may be necessary to achieve interpretable results. TROUBLESHOOTING
  b. **Analysis of T cell activation and proliferation**
    i. Perform immunohistology for CD3 and KI67 on samples co-cultured with control and target CAR-T cells as described in Box 2.
    ii. Image whole tissue sections using a confocal microscope (Fig. 5f).
    iii. Using the cell counter tool in Image J quantify the number of CD3/KI67-double positive cells and total CD3-positive cells. CRITICAL STEP Other software with cell counting functions or automated detection can be substituted for Image J.
    iv. Divide the number of CD3/KI67-double positive cells by the total number of CD3-positive cells to get the fraction of proliferating T cells. CRITICAL STEP Take measurements from multiple GBOs and regions within the same GBO and average to get a more representative measurement. TROUBLESHOOTING
  c. **Analysis of target antigen loss**
    i. Perform immunohistology for target antigen (e.g. EGFRvIII) and DAPI on samples co-cultured with control and target CAR-T cells as described in Box 2.The addition of a general tumor cell marker (e.g. SOX2 or EGFR) may help to exclude non-tumor cells within GBOs, such as immune cells, during downstream quantification. In the case of EGFRvIII, it is helpful to also analyze EGFR expression to assess CAR-T target specificity.
    ii. Image whole tissue sections using a confocal microscope (Fig. 5g).
    iii. Quantify the number of antigen-positive cells and total tumor cells using the cell counter tool in Image J.
    iv. Divide the number of antigen-positive cells by the total number of tumor cells to get the fraction of antigen-positive tumor cells. CRITICAL STEP Take measurements from multiple GBOs and regions within the same GBO and average to get a more representative measurement.
    v. Alternatively, for difficult to quantify markers (e.g. heterogeneously expressed cell surface proteins), manually outline the GBO image and measure the mean gray value of the fluorescence intensity of the target antigen in Image J as an estimate for target antigen abundance. Subtract the background fluorescence signal from each measurement for more accurate results (refer to Figure 7D in Jacob et al., 2020 for example results)^11^. CRITICAL STEP Perform immunohistology in parallel and use identical microscope settings when imaging to more accurately compare fluorescence intensity between experimental groups. TROUBLESHOOTING
  d. **Analysis of target cell killing** CRITICAL There are many methodologies for assessing cell death in GBOs, such as cleaved caspase 3 (CC3) staining, TUNEL staining, loading cells with vital dyes, chromium release, etc. but we limited this protocol to CC3 staining as an example because it allows simultaneous staining for target antigens and other cell markers to quantify the specificity of target cell killing and visualize the cytoarchitecture of GBOs during the assay. Although TUNEL staining may provide a more accurate quantification of cell death, we have not optimized this procedure. Unlike single-cell suspensions, GBOs are difficult to evenly load with vital cell dyes or chromium as a readout for cell death. These techniques may prove useful for more high-throughput analysis of cell killing if optimized.
    i. Perform immunohistology for target antigen (e.g. EGFRvIII), cleaved caspase 3 (CC3), and DAPI on samples co-cultured with control and target CAR-T cells as described in Box 2. The addition of a general tumor cell marker (e.g. SOX2 or EGFR) may help to exclude non-tumor cells within GBOs, such as immune cells, during downstream quantification. In the case of EGFRvIII, it is helpful to also analyze EGFR expression to assess CAR-T target specificity.
    ii. Image whole tissue sections using a confocal microscope (Fig 5h).
    iii. Quantify the number of antigen-positive cells, antigen-negative cells, and total tumor cells using the cell counter tool in Image J.
    iv. Divide the number of antigen and CC3-double positive cells by the total number of antigen-positive tumor cells to get the fraction of antigen-positive tumor cells that are dying. CRITICAL STEP Take measurements from multiple GBOs and regions within the same GBO and average to get a more representative measurement. CRITICAL STEP CC3 expression is lost once the cell is dead. Analyze GBOs at several time points to catch cells as they are dying.
    v. Alternatively, manually outline the GBO image and measure the mean gray value of the fluorescence intensity of CC3 in Image J as an estimate for CC3 expression. Subtract the background fluorescence signal from each measurement for more accurate results (refer to Figure 7C in Jacob et al., 2020 for example results)^11^. CRITICAL STEP Perform immunohistology in parallel and use identical microscope settings when imaging to more accurately compare experimental groups.
    vi. To assess CAR-T cell specificity, divide the number of antigen-negative/CC3-positive cells by the total number of antigen-negative tumor cells to get the fraction of antigen-negative tumor cells that are dying. TROUBLESHOOTING

## Box 1

**Processing glioblastoma tissue and GBOs for downstream analyses (Timing 2 d)**

### Procedure

1. Collect tumor pieces or GBOs into 15 mL conical tubes using a P1000 pipette with a wide bore pipette tip.
2. Carefully aspirate the medium above sunken tumor pieces or GBOs.
3. Add 10 mL of 4% formaldehyde solution to each 15 mL conical tube.
4. Incubate on a rocking platform shaker at room temperature for 30 minutes. CRITICAL STEP Proper fixation time is critical for preserving the tissue for long-term storage while maintaining the protein epitopes that primary antibodies recognize. Fixation times that exceed 30 minutes at room temperature may mask epitopes and lead to suboptimal immunohistology results.
5. Carefully aspirate the 4% formaldehyde solution above sunken tumor pieces or GBOs.
6. Wash the tumor pieces or GBOs three times with 10 mL DPBS++ by letting tumor pieces or GBOs sink to the bottom of the conical tube before carefully aspirating the supernatant.
7. Cryoprotect the tissue by adding 10 mL of 30% sucrose (wt/vol) solution to each 15 mL conical tube and incubating upright at 4°C overnight. CRITICAL STEP Tumor pieces or GBOs must sink to the bottom of the tube before proceeding to the next step. This ensures that the tissue has been fully perfused with 30% (wt/vol) sucrose.
8. Pre-label plastic cryomolds with sample identifying information using an alcohol-resistant marker.
9. Transfer tumor pieces or GBOs to plastic cryomolds using a P1000 pipette with a wide bore pipette tip. CRITICAL STEP Tumor pieces and GBOs tend to stick to the inner surface of pipette tips after incubation in 30% (wt/vol) sucrose. Keep the pipette vertical while aspirating and dispensing tissue to minimize sticking.
10. Remove excess 30% (wt/vol) sucrose solution from the cryomold by carefully aspirating using a P200 pipette. CRITICAL STEP Remove as much 30% (wt/vol) sucrose solution as possible to ensure optimal freezing without ice crystal formation.
11. Add tissue freezing medium to completely fill the plastic cryomold. CRITICAL STEP Avoid bubbles in the tissue freezing medium as they create holes after freezing which can interfere with cryo-sectioning.
12. Carefully place the tumor pieces or GBOs in the same horizontal plane in the center of the cryomold using a P200 pipette tip. This helps ensure that each cryosection contains tissue from each tumor piece or GBO. CRITICAL STEP Avoid stabbing or tearing tumor pieces or GBOs with the P200 pipette tip. Use the P200 pipette tip to manipulate the tissue freezing medium surrounding the GBOs, taking advantage of the viscosity to move them.
13. Snap freeze the cryomolds by placing in isopropanol cooled by dry ice. CAUTION Dry ice and isopropanol cooled with dry ice are very cold. Use appropriate personal protective equipment to avoid injury.
14. Dab off excess isopropanol onto a paper towel and place frozen cryomolds in a −80°C freezer for long term storage. PAUSE POINT Frozen cryomolds can be stored at −80°C indefinitely.
15. Place pre-labeled charged microscope slides and frozen cryomolds inside a tissue cryostat set to −20°C and incubate for 15 minutes. CRITICAL STEP Letting frozen cryomolds equilibrate to −20°C ensures smoother cutting.
16. Cut 25 μm sections and melt onto charged slides.
17. Dry slides for 30 minutes on a 55°C hotplate or overnight at room temperature protected from light to promote tissue adhesion to the slide.
18. Put microscope slides into a slide box and place in a −20°C freezer for long-term storage. PAUSE POINT Frozen tissue slides can be stored at −20°C for over 6 months. Consider storing at −80°C for longer time periods.
19. (Optional) To sample tissue for RNA-sequencing, place 3-5 tumor pieces or GBOs into a microcentrifuge tube using a P1000 pipette with a wide bore pipette tip, centrifuge at 500g for 5 minutes to pellet, aspirate any remaining medium, and snap freeze the microcentrifuge tubes by placing in isopropanol cooled by dry ice. Store microcentrifuge tubes containing tissue in a −80°C freezer until ready for processing. CRITICAL STEP Tumor pieces or GBOs should always be collected and analyzed in triplicates to account for variations in sampling. Individual GBOs may be sampled to analyze heterogeneity between GBOs. PAUSE POINT Frozen microcentrifuge tubes containing tissue can be stored at −80°C indefinitely.

## Box 2

**Immunofluorescence staining of glioblastoma tumors and organoids (Timing 2 d)**

### Procedure

1. Remove tissue slides from freezer storage and incubate on a hotplate set to 55°C for 30 minutes to dry. CRITICAL STEP Completely drying the tissue on a hotplate improves tissue adherence to the slide during washes and antibody incubation steps.
2. Outline tissue sections with a hydrophobic barrier pen and let dry for 5 minutes. CRITICAL STEP Avoid getting hydrophobic barrier pen ink on the tissue sections. The ink has substantial autofluorescence and its hydrophobicity prevents antibody binding.
3. Prepare immunohistology moisture chamber by adding a thin layer of ultra-pure water to the bottom of an open slide staining chamber. CRITICAL STEP This helps limit the evaporation of solution on the tissue slides during incubations when the chamber is closed.
4. Place tissue slides horizontally onto the fitted slots on the bottom of the immunohistology moisture chamber.
5. Wash tissue sections three times for 5 minutes with TBST by gently pipetting approximately 200 μL of TBST onto each tissue section and carefully aspirating the wash buffer between washes.
6. Incubate each tissue section with approximately 100 μL of blocking and permeabilization buffer for 1 hour at room temperature.
7. Aspirate blocking and permeabilization buffer and incubate each tissue section with approximately 100 μL of primary antibodies diluted in antibody dilution buffer overnight at 4°C. See Table 2 for a list of useful primary antibodies and their suggested dilutions. Dilutions for other antibodies may require optimization to achieve best signal to noise. CRITICAL STEP Only use antibody combinations with non-overlapping antibody species to avoid cross-contamination of signal. For example, use an antibody raised in mouse, rabbit, or goat together, but never two antibodies raised in the same species together. CRITICAL STEP 100 μL of antibody solution is generally enough to cover each tissue section. Adding too much solution to each tissue section may break the hydrophobic barrier and spill over to adjacent tissue sections. This will combine antibodies and render the experiment useless.
8. Aspirate primary antibody solution and wash tissue sections three times quickly and three times for 5 minutes with TBST, carefully aspirating the wash buffer between washes.
9. Incubate each tissue section with approximately 100 μL of secondary antibodies and DAPI diluted in antibody dilution buffer for 1.5 hours at room temperature. Secondary antibodies and DAPI are generally used at 1:500 dilution.
10. Aspirate secondary antibody solution and wash tissue sections three times quickly and three times for 5 minutes with TBST, carefully aspirating the wash buffer between washes.
11. Wash tissue sections once with TBS for 5 minutes to remove detergent.
12. Incubate tissue sections with 1X autofluorescence quenching solution for 1 minute. CRITICAL STEP Tumor tissue and GBOs contain blood components and lipid accumulations that have high autofluorescence that interferes with assessing true immunofluorescence signal. Incubating tissue sections with autofluorescence quenching solution is critical to block this autofluorescence. Tissue sections may turn a dark blue to black color, but antibody fluorescence is not affected.
13. Aspirate autofluorescence quenching solution and wash three times for 5 minutes with TBS. CRITICAL STEP Do not wash with buffers containing detergents as they will remove autofluorescence blocker that is bound to blood components and lipid accumulations.
14. Mount slides using approximately 50-100 μL of antifade mounting medium per slide and glass coverslips. CRITICAL STEP Avoid introducing bubbles when placing coverslips as they will interfere with imaging if located on top of tissue sections.
15. Dry slides overnight protected from light.
16. Seal the coverslips to the slides by coating the edges with clear nail polish and let dry for 1 hour.
17. Proceed to imaging with a confocal microscope (Fig. 3a) or store slides at 4°C. CRITICAL STEP While detailed discussion on microscopy technique is beyond the scope of this protocol, we emphasize optimizing microscope settings to obtain high resolution images with good signal to noise. Use a combination of secondary fluorophores with emission spectra that do not overlap substantially and avoid capturing overexposed images. For more advice on optimizing confocal microscopy refer to Jonkman, et al., 2020^52^. PAUSE POINT Immunofluorescent slides can generally be stored at 4°C for at least one month. Fluorescence signal my decrease with prolonged storage.

**Table 2.**
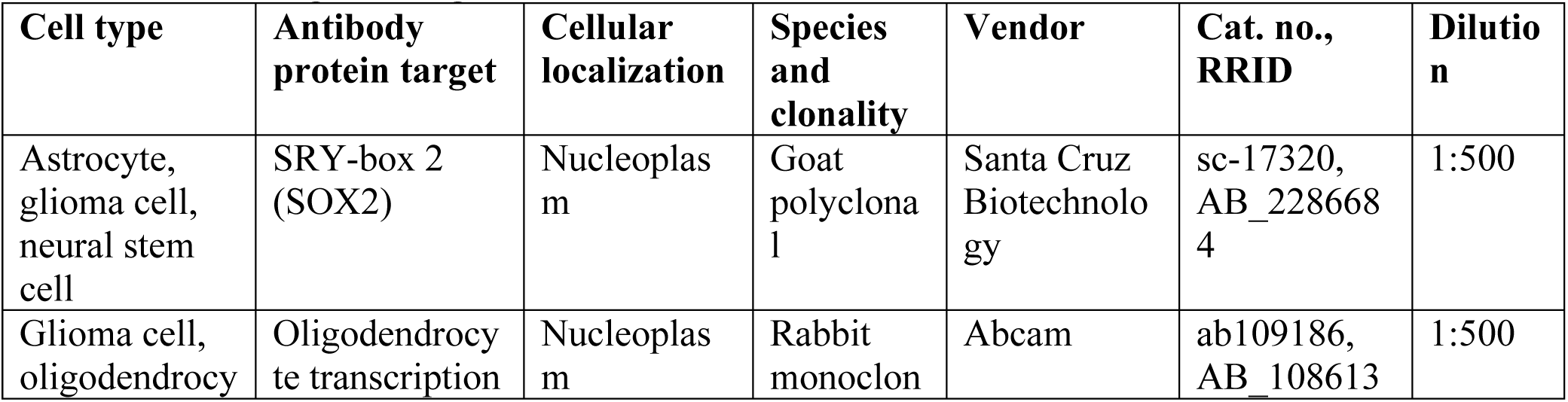
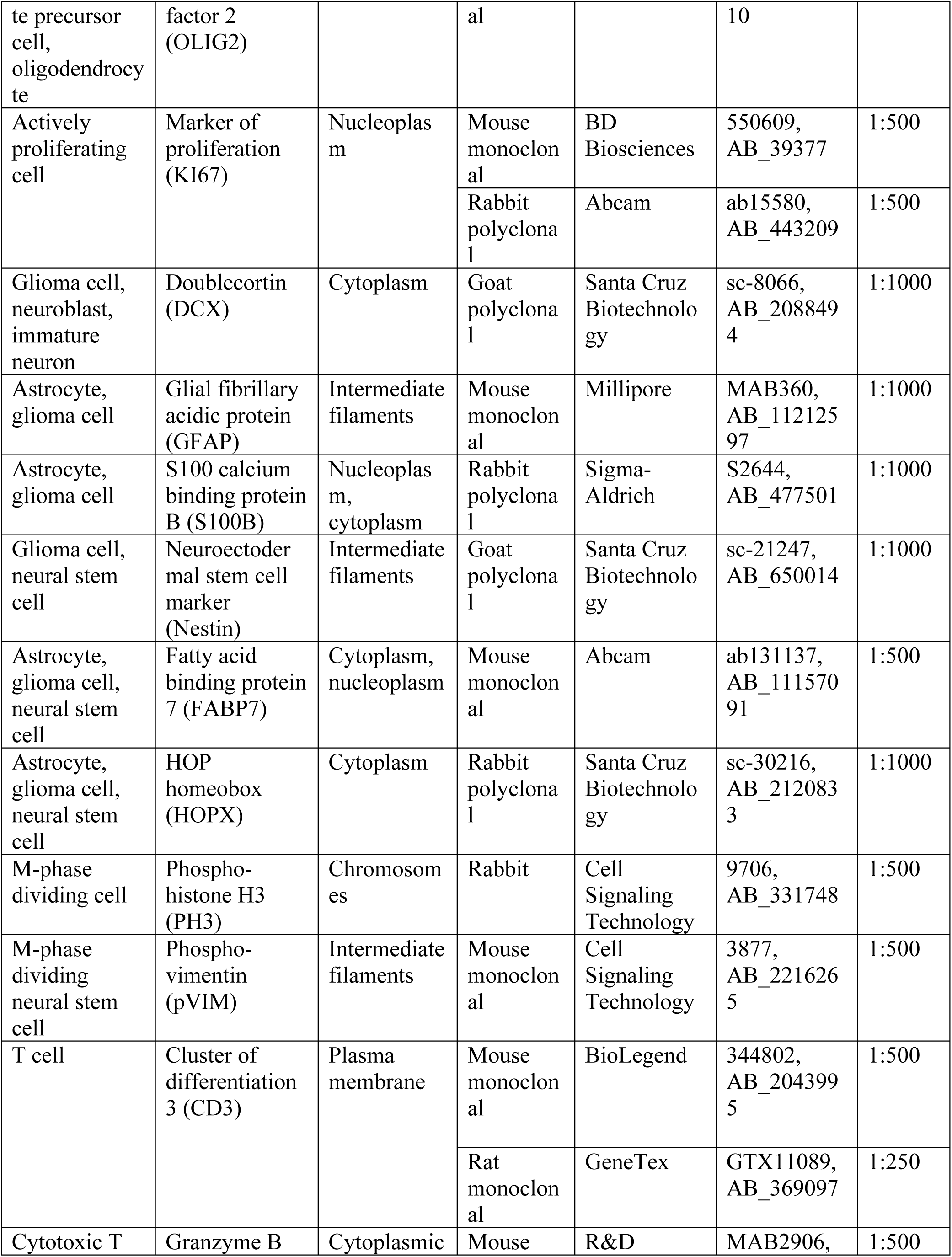
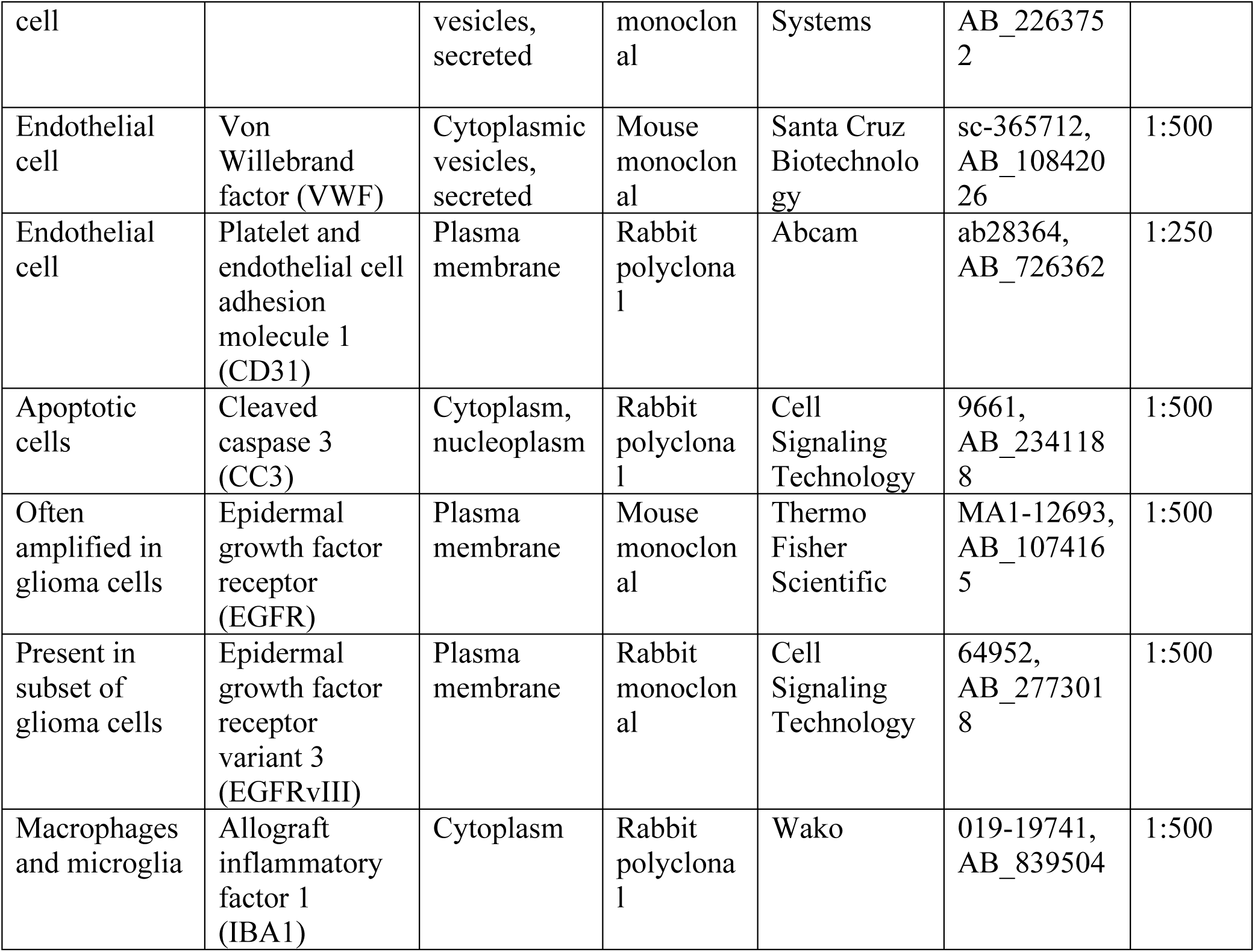
Primary antibodies used to characterize cellular heterogeneity and evaluate CAR-T activation and target killing in GBOs

## Box 3

**Detection and quantification of hypoxia gradients and actively proliferating cells in GBOs (Timing 2-3 d)**

### Procedure

1. Detection of hypoxia gradients and actively proliferating cells in GBOs is based on labeling live GBOs with pimonidazole-HCl and EdU, respectively, followed by tissue processing (Box 1) and detection with immunohistology (Box 2).
  a. **Detection of hypoxia gradients in GBOs**
    i. Prepare GBO medium containing 200 μM pimonidazole-HCl and equilibrate 4 mL per well in a 6-well plate for 15 minutes in a 37°C, 5% CO_2_, 90% relative humidity sterile incubator. CRITICAL STEP Equilibrating the medium in the incubator is critical to avoid having sudden temperature and pH changes affect the results.
    ii. Carefully add GBOs to equilibrated medium and culture for 3 hours in a 37°C, 5% CO_2_, 90% relative humidity sterile incubator.
    iii. Process GBOs for immunohistology as described in Box 1.
    iv. Perform immunofluorescence staining of tissue sections as described in Box 2 using an anti-pimonidazole primary antibody (Fig. 4a).
  b. **Detection of actively proliferating cells in GBOs** CAUTION EdU is a nucleotide analog that incorporates into newly synthesized DNA. Use appropriate personal protective equipment to avoid direct contact with EdU solutions.
    i. Prepare GBO medium containing 1 μM EdU and equilibrate 4 mL per well in a 6-well plate for 15 min in a 37°C, 5% CO_2_, 90% relative humidity sterile incubator. CRITICAL STEP Equilibrating the medium in the incubator is critical to avoid having sudden temperature and pH changes affect the results.
    ii. Carefully add GBOs to equilibrated medium and culture for 1 hour in a 37°C, 5% CO_2_, 90% relative humidity sterile incubator to label cells in the S-phase of the cell cycle.
    iii. Process GBOs for immunohistology as described in Box 1.
    iv. Perform immunofluorescence staining of tissue sections as described in Box 2 using the instructions for the Click-iT EdU detection kit (Fig. c-d).
    v. (Optional) Perform immunofluorescence staining for phospho-vimentin (pVIM) and phospho-histone H3 (PH3) as described in Box 2 to detect and analyze the morphology of tumor cells in mitosis (Fig. 4e).
2. Quantification of hypoxia gradients and actively proliferating cells is performed after imaging with a confocal microscope using image analysis software, such as Image J.
  a. **Quantification of hypoxia gradients**
    i. Image whole GBO tissue sections using a confocal microscope (Fig. 4a). CRITICAL STEP Take images using an appropriate laser power that does not result in overexposed regions. Quantification of fluorescent signal intensity is not accurate when regions are overexposed.
    ii. Using Image J, draw a line from the edge of the GBO to the center.
    iii. Measure the mean gray value of the Hypoxyprobe fluorescent signal as a function of distance using the plot profile function in ImageJ (Fig. 4b). CRITICAL STEP Take multiple measurements starting from different edges of the same GBO and multiple GBOs treated in parallel and average to get a more representative measurement. CRITICAL STEP Other software capable of measuring fluorescent intensity can be substituted for Image J.
    iv. (Optional) Measure the distance of each proliferating cell (labeled by KI67 or EdU) from the surface of the GBO and plot the abundance of KI67^+^ cells as a function of distance (Fig. 4b).
  b. **Quantification of actively proliferating cells**
    i. Image whole GBO tissue sections using a confocal microscope (Fig. 4c).
    ii. Using the cell counter tool in Image J, count the total number of nuclei using the DAPI immunofluorescence. CRITICAL STEP Other software with nuclei counting functions or automated nuclei detection can be substituted for Image J.
    iii. Count the number of actively proliferating nuclei using the EdU immunofluorescence using the cell counter tool.
    iv. Divide the number of EdU-positive cells by the total number of nuclei to get the fraction of actively proliferating cells. CRITICAL STEP Take measurements from multiple GBOs and regions within the same GBO and average to get a more representative measurement.
    v. EdU detection can be combined with marker immunohistology to quantify the proliferation rates of different tumor cell populations^11^ (Fig. 4d). For this, use the total number of marker-positive cells as your denominator and EdU/marker-double positive cells as the numerator.

## Troubleshooting

Troubleshooting advice can be found in Table 3.

**Table 3.**
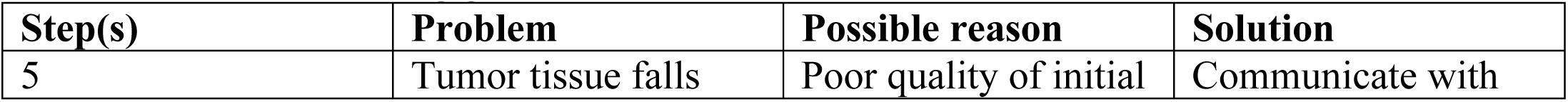
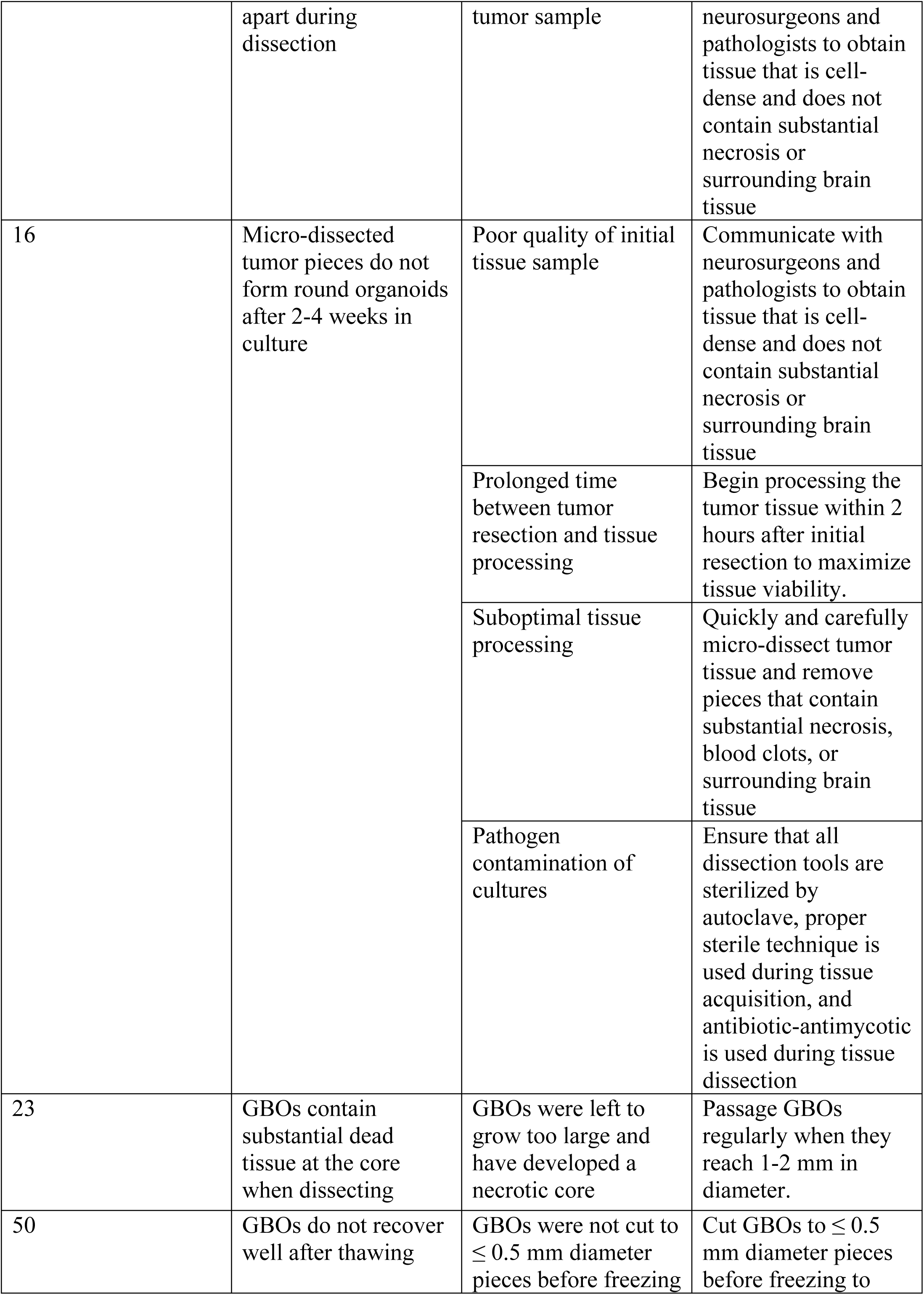
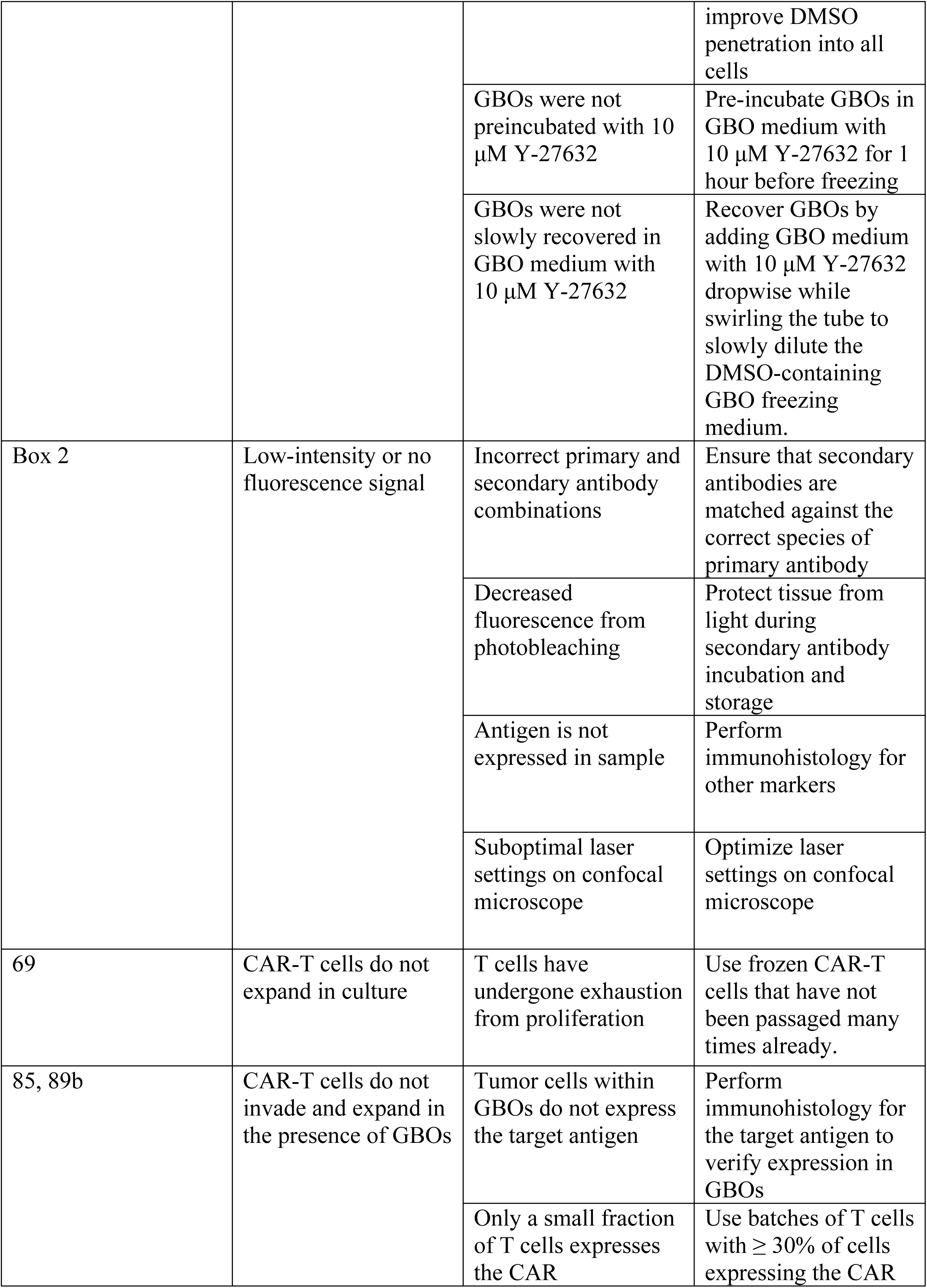
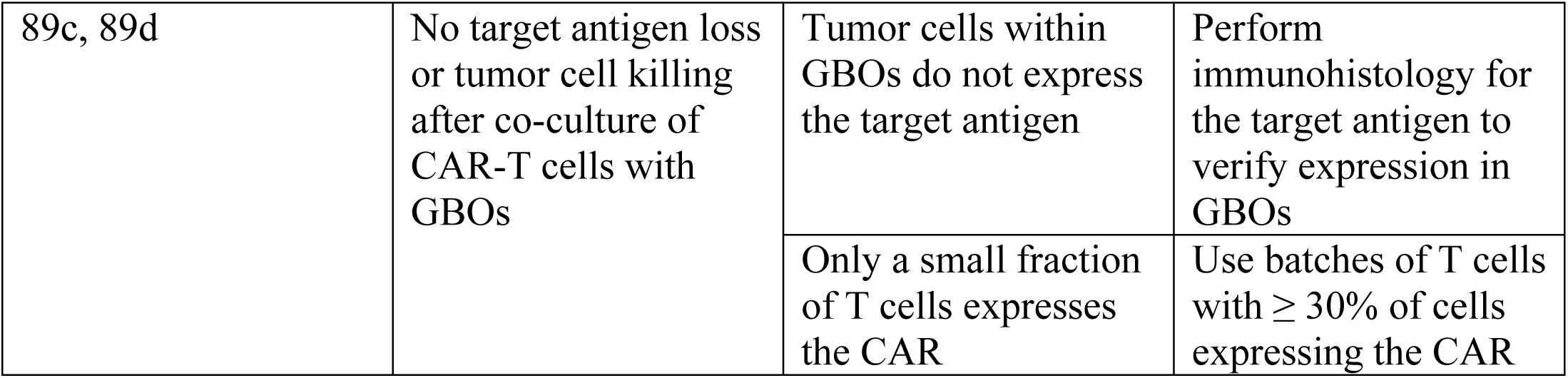
Troubleshooting guide

## Timing

- Steps 1-19, establishment of glioblastoma organoid culture from resected tumor tissue: 2-4 h for tissue processing and 2-4 w for organoid establishment
- Steps 20-26, passaging glioblastoma organoids: 1-2 h
- Steps 27-39, cryopreservation of glioblastoma organoids: 2-3 h
- Steps 40-50, thawing glioblastoma organoids: 1 h
- Steps 51-69, preparation of CAR-T cells for co-culture: 1 h for thawing, 3 days for recovery and expansion
- Steps 70-88, co-culture of GBOs with CAR-T cells: 1-5 d
- Step 89, downstream assays: variable
- Box 1, processing glioblastoma tissue and GBOs for downstream analyses: 2 d
- Box 2, immunofluorescence staining of glioblastoma tumors and organoids: 2 d
- Box 3, detection and quantification of hypoxia gradients and actively proliferating cells in GBOs: 2-3 d

## Anticipated results

The protocol described here enables the user to generate GBOs from primary and recurrent/residual resected glioblastoma tumor tissue and co-culture GBOs with CAR-T cells to evaluate cell killing and target specificity. We have generated GBOs from 95.7% of primary tumors and 75.0% of recurrent/residual tumors for an overall success rate of 91.4% (Fig. 1b). We have had success making GBOs from tumors containing a wide range of mutations commonly found in glioblastoma (Fig. 1c). Owing to the rapid generation of GBOs in 2-4 weeks following surgery, we have an average time to establish GBO biobank of 27 days (Fig. 1d). GBOs grow at different rates and have different cell numbers depending on the size and density of the tumor cells within them (Fig. 1e-f), Although the results will vary depending on the tumor-specific growth characteristics of each sample and the CAR-T cells used, there are general results one should expect for GBO generation and CAR-T cell co-culture.

## Evaluation of successful GBO generation and cryopreservation

Successful GBO generation can be assessed visually by bright-field and confocal microscopy. GBOs should always be compared to their primary tumors to assess recapitulation of important characteristics, including proliferation rates, marker immunohistology, transcriptional signatures, and genomic mutations. Be sure to evaluate at least 5 different GBOs at multiple timepoints to assess and account for sample heterogeneity. Optimal tumor tissue is resected *en bloc* from cell-dense regions of the tumor with minimal necrosis, hemorrhage, or surrounding brain tissue (Fig. 2c). Micro-dissected tumor pieces should round up and form organoids with a fairly homogenous off-white color with sharp borders and few floating single cells within 2-4 weeks of culture (Fig. 2g). GBOs should continue growing for at least 6 months with proper culture and passaging by sequential bisection (Fig. 2k-l). Although volumetric and weight measurements are difficult, GBO growth can be assessed by measuring the increase in 2D area over time (Fig. 1e). GBOs should retain the cellular heterogeneity and proliferation rates of parent tumors as evidenced by immunohistology for various markers for glioma stem, progenitor, and differentiated cells (Fig. 3a). Successful GBOs should also recapitulate the transcriptomes and mutations of their parent tumors (refer to data in Jacob et al., 2020)^11^. GBOs should retain immune cells, such as IBA1^+^ tumor-associated macrophages/microglia for at least 6 weeks, although their numbers will vary between tumors and always decrease with time due to death and dilution by proliferative tumor cells (Fig. 3b-c). GBOs will often also retain endothelial cells and vasculature, although their abundance may vary greatly between tumor samples and their functionality has not been well characterized (Fig. 3d-e). Unsuccessful GBO generation is most likely a product of poor-quality resected tumor samples that contain few viable tumor cells and abundant necrosis. Unsuccessful cultures are often composed of tumor sparse tissue (Fig. 2f) and/or necrotic pieces (red arrow; Fig. 2i). Failed GBO cultures are noticeable as pieces that fail to round-up and grow after 2-4 weeks (Fig. 2j).

Successful cryopreservation of GBOs is evaluated by recovery after thawing. Thawed GBOs should appear as solid pieces that begin to round-up and grow within 2 weeks in culture (Fig. 2m). Thawed GBOs may have fuzzy borders with some dead floating cells immediately after thawing, but the borders should become sharp as they proliferate under orbital shaking conditions. Growth of thawed GBOs can be assessed by measuring the increase in 2D area over time, which will increase on average from >1 to 3-fold over 4 weeks depending on tumor-specific characteristics and the quality of cryopreservation (Fig. 1e). Successfully cryopreserved and thawed GBOs should be cell-dense and retain similar marker immunohistology to their parent tumors and GBOs before cryopreservation (Fig. 3a) with few apoptotic cells (Refer to Figure S1A-B in Jacob et al., 2020)^11^. We recommend testing GBO recovery after cryopreservation to assess the quality of the biobank for each sample.

## Evaluation of hypoxia gradients and actively proliferating cells in GBOs

Large GBOs (> 2 mm in diameter) will contain increased hypoxia towards the core with a higher density of proliferating cells towards the GBO surface (Fig. 4a-b). Pulsing GBOs with EdU for 1 hour will label cells in S-phase of the cell cycle (Fig. 4c-d). EdU will often co-localize with glioma stem and progenitor cell markers, such as NESTIN, HOPX, OLIG2, and DCX (Fig. 4d). Immunohistology for phospho-vimentin and phospho-histone H3 will mark cells in M-phase of the cell cycle and reveal diverse morphologies of actively dividing cells which often resemble cells of the developing human brain (Fig. 4e), such as radial glia which have been described in glioblastomas^53,54^.

## Evaluation of successful GBO co-culture with CAR-T cells

For optimal results, verify expression of CAR-T cell target antigen in parent tumors and GBOs (Fig. 5b) and include control CAR-T cells which recognize an antigen, such as CD19, not expressed by parent tumors or GBOs. By bright-field microscopy, co-culture of GBOs with CAR-T cells should show variable T cell proliferation and invasion into GBOs depending on the presence of target antigen; T cells will proliferate in clusters around GBOs that express the target antigen (Fig. 5c-d). CAR-T cells that do not recognize an antigen within GBOs should not substantially increase their proliferation rate and kill any tumor cells. CAR-T cells that recognize an antigen on tumor cells within GBOs will invade, proliferate, and kill tumor cells as evidenced by loss of target antigen and an increase in CC3 immunoreactivity (Figs. 5g-h). CAR-T cells that recognize antigen will proliferate and be found predominately at the periphery of GBOs in clusters as CD3/KI67 double-positive cells (Fig. 5f). The length of co-culture needed to observe killing will depend on many factors, including the abundance of target antigen-expressing tumor cells, the binding kinetics of the CAR, and variations between different GBO samples. Increased release of cytokines (e.g. IL-2, IFNγ, TNFα) from activated T cells should occur within 24-72 hours after co-culture in the presence of target antigen (Fig. 5e)^11^. In our experiments with 1:10 CAR-T cells to tumor cells, we observed a spike in IL-2 levels after 24 hours, a peak in TNFα levels at 24 hours with sustained secretion up to 72 hours, and continuously increasing levels of IFNγ up to 72 hours after co-culture. The timeline and levels of cytokines may vary depending on antigen abundance and CAR-T cell ratio, health, and specificity.

## Reporting Summary

Further information on research design is available in the Nature Research Reporting Summary linked to this article.

## Data Availability

The data that provide examples of the results that can be generated with this protocol are available from the corresponding authors on reasonable request.

## Acknowledgements

We thank all patients who generously donated tissue. We thank members of the Ming and Song laboratories for comments and suggestions; B. Temsamrit and E. LaNoce for technical support; J. Schnoll for lab coordination; and D. O’Rourke, M. Nasrallah, and others in the Departments of Neurosurgery and Pathology at the Hospital of the University of Pennsylvania for assistance with tissue acquisition. This work was supported by Glioblastoma Translational Center of Excellence at the Abramson Cancer Center at University of Pennsylvania, grants from National Institutes of Health (R37NS047344 and R35NS116843 to H.S., R35NS097370 and U19AI131130 to G-l.M.), and the Sheldon G. Adelson Medical Research Foundation (to G-l.M.).

## Author contributions

F.J. developed the procedures to generate, culture, and biobank glioblastoma organoids and to co-culture glioblastoma organoids with CAR-T cells and produced the sample data. F.J., G.-l.M., and H.S. conceived the project and wrote the manuscript.

## Competing interests

The authors declare no competing interests.

## List of Supplementary Information

Movie 1. Micro-dissection of resected tumor tissue.

Supplemental Methods

## Notes

### Competing Interest Statement

The authors have declared no competing interest.

### Summary of Updates

Revised version that incorporates suggestions from peer-review.

