## Supplemental Methods for "Generation and biobanking of patient-derived glioblastoma organoids and their application in CAR-T cell testing"

### Additional Materials

#### Reagents

- 2173 (EGFRvIII) CAR construct (Johnson et al., 2015)[1]
- Biotinylated Goat Anti-Human IgG, F(ab')2 fragment specific (Jackson ImmunoResearch Labs, cat. no. 109-066-006, RRID: AB_2337634)
- Biotinylated Goat Anti-Mouse IgG, F(ab')2 fragment specific (Jackson ImmunoResearch Labs, cat. no. 115-066-072, RRID: AB_2338583)
- CD19 CAR construct (Porter et al., 2011)[2]
- Donor leukapheresis products
- Dynabeads Human T-Activator CD3/CD28 (ThermoFisher Scientific, cat. no. 11131D)
- Fetal bovine serum (FBS; GE Life Sciences, cat. no. SH30071.03)
- HEPES, 1 M (ThermoFisher Scientific, cat. no. 15630080)
- R-Phycoerythrin Streptavidin (Jackson ImmunoResearch Labs, cat. no. 016-110-084, RRID: AB_2337240)
- RPMI 1640 medium (ThermoFisher Scientific, cat. no. 11875093)

#### Equipment

- Multisizer 3 Coulter Counter (Beckman Coulter)
- LSR II flow cytometer (BD Biosciences)

#### Software

- FlowJo (https://www.flowjo.com/solutions/flowjo, RRID:SCR_008520)

### Procedures

#### Generation of CAR-T cells from donor leukapheresis products[1,3]

Peripheral T cells were isolated from leukapheresis products obtained from de-identified healthy donors under a protocol approved by the University of Pennsylvania's Institutional Review Board. Isolated T cells were stimulated with Dynabeads Human T-Activator CD3/CD28 at a bead to cell ratio of 3:1. T cells were cultured in RPMI 1640 medium supplemented with 10% fetal bovine serum, 20 mM HEPES, and PenStrep and medium was replaced every 48 hours. The end of the stimulation was determined by a decrease in log-phase growth and reduced mean lymphocytic volume to 300-330 fl as measured on a Coulter Multisizer. This was usually reached 10 days after stimulation. The T cells were transduced with lentiviruses containing the CD19 or 2173 (EGFRvIII) CAR transgenes. To determine the percentage of T cells expressing the desired CAR transgene, a $1 \times 10^6$ cells/ml T cell sample in PBS containing 0.2% BSA was incubated with either biotinylated goat anti-human antibody for the 2173 CAR (human scFv) or biotinylated goat ant-mouse antibody for the CD19 CAR (mouse scFv) for 30 minutes at 4°C. The T cells were washed twice with DPBS++ containing 0.2% BSA and secondary detection was carried out by the addition of streptavidin-coupled PE for 30 minutes at 4°C. T cells were washed twice with DPBS++ containing 0.2% BSA and resuspended in DPBS++ with 2% formaldehyde. Fluorescence was assessed using a BD LSR II flow cytometer, and data were analyzed with FlowJo software. For this present study, T cells transduced with CD19 and 2173BBz CARs were 30% and 32% $CAR^+$, respectively.
